# Distinct extracellular-matrix remodeling events precede symptoms of inflammation

**DOI:** 10.1101/665653

**Authors:** Elee Shimshoni, Idan Adir, Ran Afik, Inna Solomonov, Anjana Shenoy, Miri Adler, Luca Puricelli, Veronica Ghini, Odelia Mouhadeb, Nathan Gluck, Sigal Fishman, Lael Werner, Dror S. Shouval, Chen Varol, Alessandro Podestà, Paola Turano, Tamar Geiger, Paolo Milani, Claudio Luchinat, Uri Alon, Irit Sagi

## Abstract

Identification of early processes leading to complex tissue pathologies, such as inflammatory bowel diseases, poses a major scientific and clinical challenge that is imperative for improved diagnosis and treatment. Most studies of inflammation onset focus on cellular processes and signaling molecules, while overlooking the environment in which they take place, the continuously remodeled extracellular matrix. In this study, we used colitis models for investigating extracellular-matrix dynamics during disease onset, while treating the matrix as a complete and defined entity. Through the analysis of matrix structure, stiffness and composition, we unexpectedly revealed that even prior to the first clinical symptoms, the colon displays its own unique extracellular-matrix signature and found specific markers of clinical potential, which were also validated in human subjects. We also show that the emergence of this pre-symptomatic matrix is mediated by sub-clinical infiltration of neutrophils and monocytes bearing remodeling enzymes. Remarkably, whether the inflammation is chronic or acute, its matrix signature converges at pre-symptomatic states. We suggest that the existence of a pre-symptomatic extracellular-matrix is general and relevant to a wide range of diseases.

## Introduction

The majority of research efforts on complex diseases are invested in studying cellular processes and cellular signaling molecules, e.g., cytokines, chemokines and growth factors, which often represent only a subset of factors responsible for tissue pathology. All of these processes and molecular signals are embedded within the extracellular matrix (ECM), which comprises a large portion of the tissue. Due to the intimate cell-matrix relationship, aberrations in the ECM hold the potential to induce pathological cellular behavior[1–3]. Recent studies suggest that dysregulated production of ECM modulators plays key roles in inflammation[4]. Along these lines, we hypothesized that the ECM can serve as an early reporter for multifactorial inflammatory diseases resulting in tissue damage. We thus set to investigate our hypothesis in inflammatory disease models in which tissue damage and ECM remodeling are known to occur[5].

Inflammatory bowel diseases (IBD) is an umbrella term for idiopathic diseases in the digestive tract, namely Crohn’s disease (CD) and Ulcerative Colitis (UC). Early clinical intervention is key for preventing the cascading of chronic inflammation into irreparable tissue damage. However, this is challenging since, similarly to many idiopathic complex disorders, diagnosis is determined according to symptoms that appear well after molecular pathological processes are already taking place.

A number of studies over the past two decades have found ECM remodeling enzymes to be upregulated in human IBD. In a complimentary manner, the key role of ECM composition and remodeling in the onset, progression and severity of IBD has been implicated in various rodent models[6–14]. We have previously provided preliminary evidence for the structural deformation taking place in the colonic ECM of IBD patients[5]. Accordingly, we set out to explore early involvement of ECM remodeling in the disease, before any gross signs of inflammation are detected.

In this study, we monitored ECM dynamics during intestinal inflammation by utilizing two common murine models for IBD – the acute dextran sodium sulfate (DSS)-induced colitis model[15], and the chronic piroxicam-accelerated colitis (PAC) model in interleukin(IL)-10^-/-^ mice[16], and then validating them using available human specimens from healthy individuals and IBD patients. Through the analysis of matrix structure, stiffness and composition, we unexpectedly revealed a pre-symptomatic state with its own unique ECM signature and found markers of clinical potential. In addition, we show that this signature emerges, at least in part, by the increased activity of remodeling enzymes originating from infiltrating monocytes and neutrophils and the epithelium. Our integrated analysis highlights the ECM as a central component of early inflammatory processes and its potential in serving a prognostic biomarker.

## Results

### Characterization of murine colitis models for mapping ECM properties and dynamics

To probe the properties of native ECM during disease-associated transitions we adopted established murine models for IBD. Noteworthy, the study of IBD is almost exclusively immunological and cell-centric, with only limited study of the ECM and lack of information regarding its material properties, since it is usually relegated to background or contextual consideration. We have challenged this bystander role for the ECM[5].

To begin, we use a common murine model for IBD – the DSS-induced colitis model[15], which, with the appropriate calibration, results in a well-defined time-frame of acute colitis development in wild-type (WT) mice (**Fig. 1A**, see Methods for details). Disease progression was monitored by body weight measurement, endoscopic evaluation and histological analysis of the colon, in accordance with commonly used clinical diagnostic methods[17] at different time points – day 0 (healthy WT), day 4 and day 10 (**Fig. 1** and **Fig. S1,** see Methods section for details on endoscopic scoring). Colon inflammation under these conditions peaks on day 10, as detected by endoscopy and histology (**Fig. S1** and **Fig. 1B-D**). Importantly, while clinical symptoms begin to appear after day 5, on day 4 most animals (over 90%) are defined as “healthy” (**Fig. 1C**), since they do not lose weight (**Fig. 1E**), appear healthy by endoscopic evaluation, and do not display inflammation that is consistently detectable via histopathological analysis (**Fig. 1B** and **1D**). Therefore, we regard day 4 as the “pre-symptomatic” state for analysis of ECM dynamics before the emergence of clinical symptoms.

**Figure 1.**
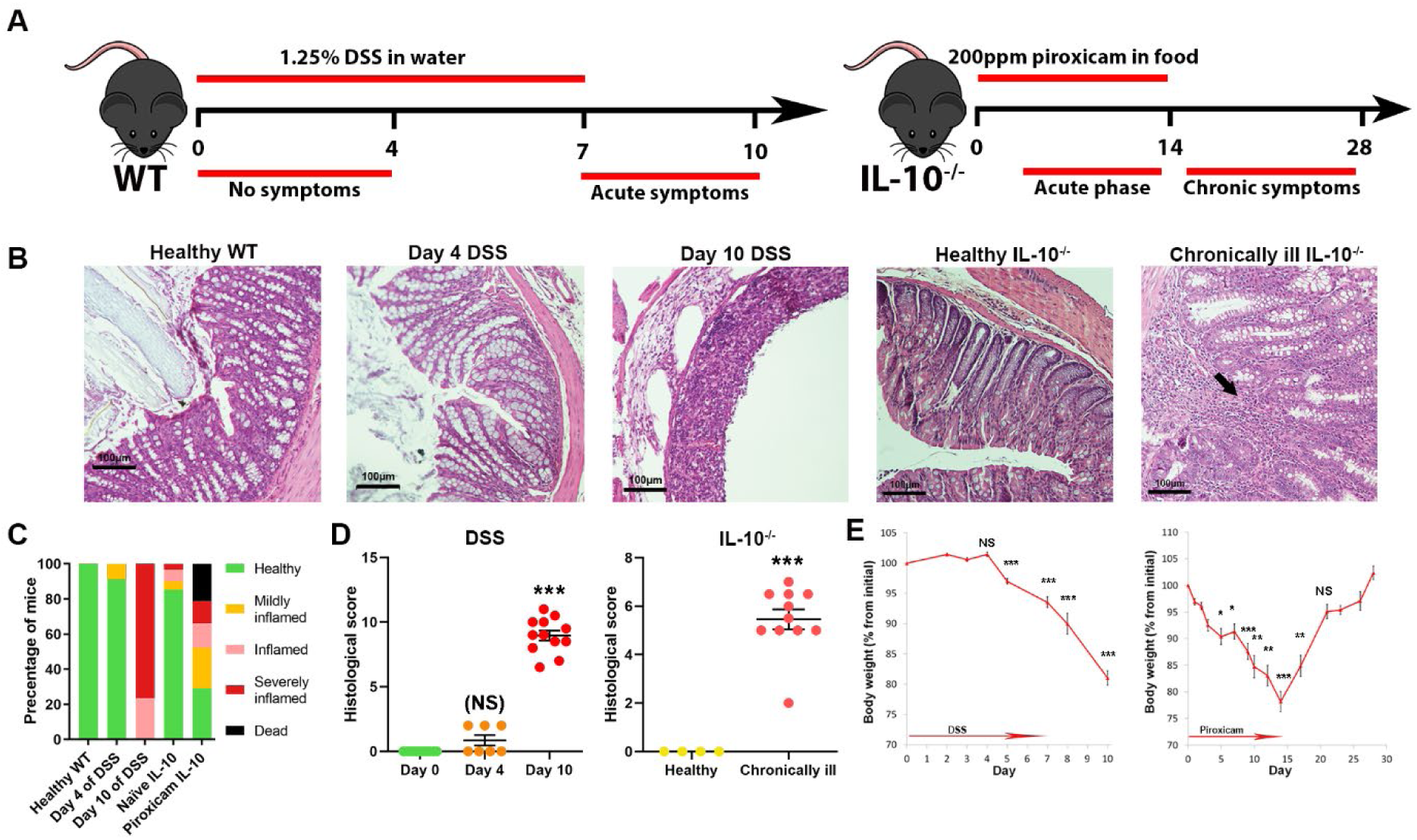
Definition of the disease course and tissue states in the two murine colitis models. (A) Acute model – DSS-induced colitis. Clinical and endoscopic symptoms are apparent from day 7 and peak at day 10. Chronic model – PAC IL-10^-/-^ model. Clinical and endoscopic symptoms develop over the course of 14 days, and the chronic inflammatory state is persistent per mouse. Chronically ill IL-10^-/-^ mice were harvested at least two weeks following piroxicam discontinuation. (B) H&E-stained colonic sections of mice from the two models at the indicated states. Note that immune cell infiltration and mucosal damage is not evident on day 4 of the acute model, but is substantial on day 10. In the PAC IL-10^-/-^ model large amounts of immune cell infiltrate is observed in the mucosa (indicated by arrow), but is absent in the healthy IL-10^-/-^ mouse. (C) Quantification of the percentage of mice in each clinical category according to endoscopic evaluation (0-4: “healthy”; 5-7 “mildly inflamed”; 8-11 “inflamed”; and 12-15 “severely inflamed”, see **Fig. S1** and Methods for details) at the indicated time points or states of both models (n>30). Note that some IL-10^-/-^ mice develop spontaneous inflammation without piroxicam exposure, and that some do not develop inflammatory symptoms following exposure. Only mice evaluated as “healthy” were used for further analysis as the day 4 of DSS or healthy IL-10^-/-^ states. (D) Histological scoring of H&E stained sections, such as the ones shown in B. Details on scoring are available in the Methods. In the DSS model, a significant increase in histopathological score was only observed on day 10 and not on day 4 (NS=not significant). Chronically ill IL-10^-/-^ mice display a significant increase in histopathological score. The plot displays the scores of individual animals and the bars represent the state’s mean±s.e.m. Statistical significance was determined using a student’s t-test with a Bonferroni correction for two comparisons to healthy WT for the DSS model (NS: P> 0.025, *** P<0.0005) and with no correction, compared to healthy IL-10^-/-^ in the PAC IL-10^-/-^ model (*** P<0.001). (E) Mouse body weight changes over the course of the two models, presented as percentage from body weight on day 0. Significant weight loss in the DSS model appears from day 5 compared to day 2. Number of animals: n(2)= 85, n(4)=55, n(5)=36, n(7)=41, n(8)=10, n(10)=52; Bonferroni for five comparisons: ***P<0.0002. Significant weight loss appears in the IL-10^-/-^ mice from day 5 compared to day 1. Number of animals: n(1)= 8, n(5)=22, n(7)=7, n(9)=7, n(10)=13, n(12)=19, n(14)=10, n(17)=10, n(21)=17; Bonferroni correction for eight comparisons: *P<0.00625, **P<0.00125, ***P<1.25X10^-4^. Note that mice regain their weight as they reach the chronic phase of colitis.

Since our goal was to use a range of ECM states along the health-disease axis in intestinal inflammation, we also examined C57BL/6 IL-10^-/-^ mice (“IL-10”) that spontaneously develop chronic colon inflammation[18]. In these mice, inflammation is not chemically induced, but rather is thought to result from immune dysregulation, and phenotypically, they more closely resemble the clinical manifestation of chronic UC and CD patients and mimic a monogenic form of IBD[16, 19] (**Fig. 1A**, see Methods for details). IL-10 mice develop IBD-like symptoms, which can be detected and scored using colonoscopy, similarly to the well-established DSS-induced model (**Fig. S1**). Histopathology in this model shows prominent immune cell infiltration into the mucosa and has a different scoring system than DSS (see Methods for details) (**Fig. 1B**). Importantly, healthy naïve IL-10 mice do not display histopathological parameters of inflammation, while ill mice have a high histopathological score of 5.5 out of 8 on average (**Fig. 1D**). The penetrance of disease in this model is different than in the DSS-induced colitis, as just over 70% of the mice develop colitis at varying degrees, with a ∼20% mortality rate (**Fig. 1C**). For comparing acute (day 10 of DSS) and chronic inflammation, we selected a stable chronic-inflammation time point for further investigation (day 28), when mice appear to be at, or surpass, their initial weight before treatment (**Fig. 1E**). For comparison to the pre-symptomatic day 4 time point in the DSS model, we chose to use naïve eight to twelve-week-old IL-10 mice that were deemed “healthy” by endoscopic evaluation. Taken together, pre-symptomatic states and clear disease states were defined in two animal models: one for transient acute colitis (DSS) and the other for persistent chronic colitis (IL-10).

### ECM morphological changes precede inflammatory symptoms

Having established pre-disease and disease state conditions, we sought to analyze the dynamics of ECM morphology by direct visualization. To do this, we first utilized second-harmonic generation (SHG) microscopy, which allows visualization of fibrillar collagen, without labeling, in native tissues. More detailed and high-resolution inspection of the spatial organization of the ECM was achieved using scanning electron microscopy (SEM) on decellularized colon tissue from the same mice.

Integration of SHG and EM analyses goes beyond the histological level, providing a detailed architectural depiction of the ECM in each tissue state namely, WT, pre-symptomatic, acute and chronic inflammation. Remarkably, marked structural and morphological changes in ECM could be observed already before any clinical or histological evidence of inflammation (**Fig. 1**). Below we describe each state and its unique architectural ECM signature.

#### Healthy WT colonic ECM architecture

SHG microscopy revealed that in the colon of the healthy WT mouse, the collagen (**Fig. 2A**) uniformly circumscribes the colonic crypts with little to no signal of fibrillar collagen between crypt borders (borders are pointed out by arrows in **Fig. S2A**). The tissue architecture in this state is well-organized, with a homogeneous crypt diameter (mean=53.7µm, average STD=10.8µm) and crypt wall thickness (mean=12.8µm, average STD=6.1µm) (**Fig. 2C**). High-resolution SEM reveals that healthy crypts are uniformly covered by a dense mesh layer, which is most likely the basement membrane, according to its morphology[20] (**Fig. 2B**).

**Figure 2.**
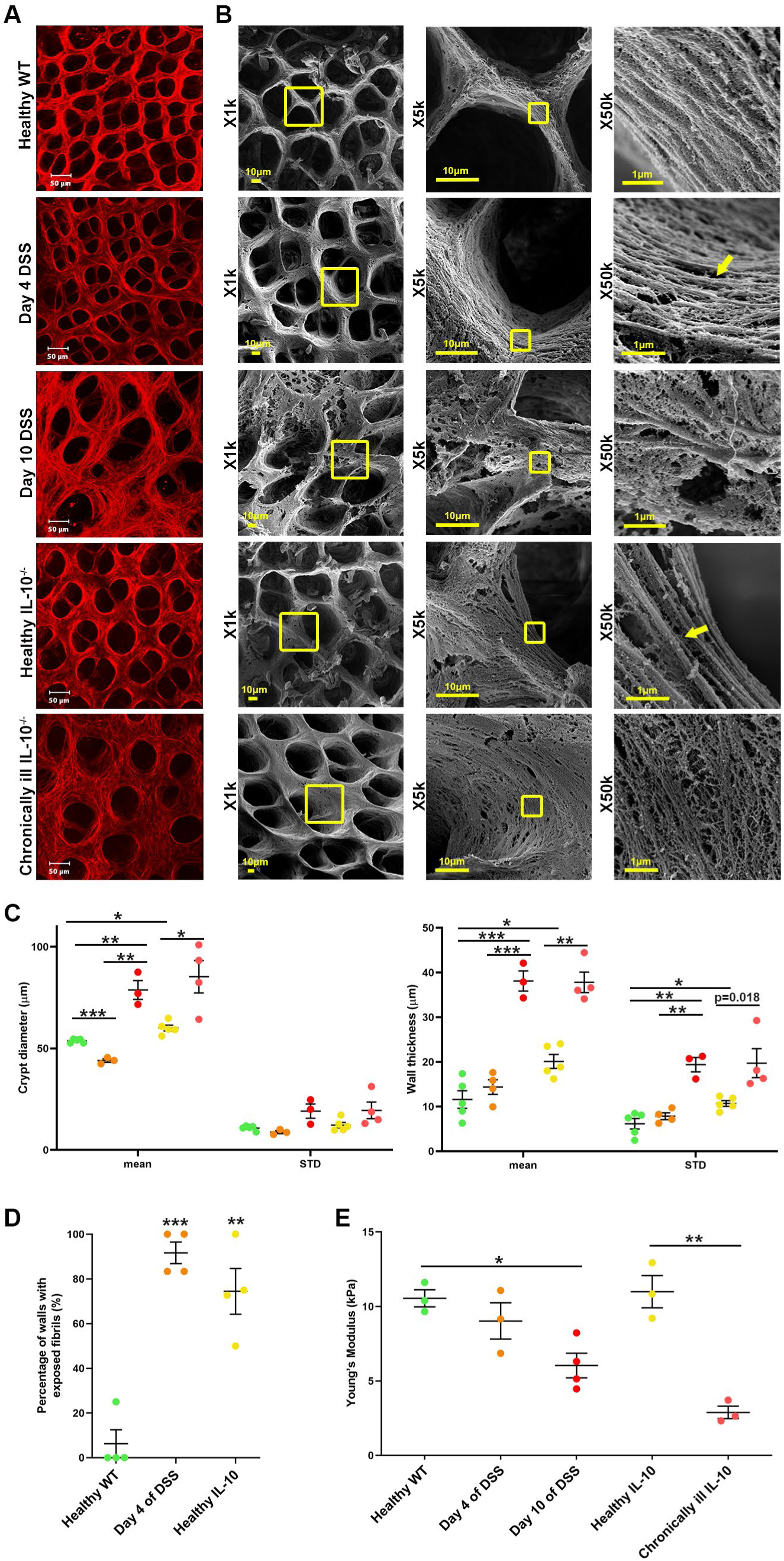
The ECM suffers structural damage and softening during colitis. (A) Second-harmonic imaging of mouse colon on the indicated time points over the course of the disease, corresponding to distinct states: healthy WT, pre-symptomatic (day 4 of DSS and healthy IL-10^-/-^), acute inflammation (day 10 of DSS) and chronic inflammation (Chronically ill IL-10^-/-^). Vast ECM structural changes occur in acute inflammation, as indicated in the comparison between day 0 and day 10. Remarkably, some reorganization of the ECM is apparent from day 4, before clinical symptoms are evident. Healthy IL-10^-/-^ display different ECM structure than that of WT mice, with ECM condensation similar to that of day 4. Chronic illness leads to overall maintained crypt architecture, but with loosely packed ECM. (B) SEM images of decellularized colonic ECM at the corresponding states, at three different magnifications. Note the damage and crypt size heterogeneity apparent on day 10 compared to Healthy WT colon. Also note, that the ECM under chronic illness is characterized by perforated crypt walls with thin fibrils. Most remarkably, the two pre-symptomatic states show a common feature of exposure of fibrillar ECM proteins on the crypt walls, as can be observed in the X50k magnification (examples are indicated by arrows). (C) Measurement of crypt diameter and crypt-wall thickness in second-harmonic images on all five states – healthy WT (green), DSS day 4 (orange), DSS day 10 (red), healthy IL-10^-/-^ (yellow) and ill IL-10^-/-^ (pink). In oval crypts, the largest diameter was always chosen, and the shortest distance was taken for wall thickness measurements. Note that each state has a different mean diameter and wall thickness. Changes in the standard deviations (STDs) of these measures per animal are also apparent, demonstrating their heterogeneity. Both of these measures indicate the architectural changes taking place in the ECM. Statistical significance was determined using a t-test with a Bonferroni correction for five comparisons, and is indicated by asterisks. *P<0.01, **P<0.002, ***P<0.0002. The bars represent the mean±s.e.m of the state, while each dot corresponds to the mean/standard deviation of one animal, which is based on the analysis of 2-8 440µmX440µm images capturing n>30 crypts per animal. One outlier was removed in the day 4 group. (D) The relative frequency of exposed fibrils on crypt walls in the SEM images, indicating a leaky ECM underlying the epithelium, in Healthy WT vs. Day 4 of DSS and Healthy IL-10^-/-^ samples. **P<0.005, ***P<0.0005, n=4 different samples per state, based on analysis of n>14 crypts. (E) ECM softening during colitis progression: individual dots represent the properly weighted median values of the Young’s modulus for each animal and the bars represent the average median±STD, corresponding to all distributions represented in **Fig. S3**. There is a significant reduction in ECM rigidity between the healthy WT and acutely inflamed samples as denoted by the asterisks (according to a t-test with Bonferroni correction for three comparisons, *P<0.017). The differences between the pairs HC-D4 and D4-D10 are not statistically significant (P=0.30 and P=0.09, respectively); nevertheless, pre-symptomatic ECMs (D4) show an intermediate behavior with respect to the healthy and acutely inflamed cases. Also, ECM rigidity significantly reduces as IL-10^-/-^ mice become chronically ill, as denoted by the asterisks (**P<0.01).

#### Acute colitis ECM architecture

Both imaging modalities reveal that acute inflammation (day 10 of DSS) has a profound effect on ECM structure. This state presents with destruction of tissue architecture (**Fig. 2A**), altered and heterogeneous crypt (mean=78.7µm, average STD=19.1µm) and wall size (mean=38.1µm, average STD=19.4µm) (**Fig. 2C**), and disrupted ECM morphology (**Fig. 2B** and **Fig, S2B**), as well as regions containing the high-intensity fibrillar structures highlighted in **Fig. S2A**. These findings indicate that acute inflammation is characterized by regional ECM degradation and build up.

#### Chronic colitis ECM architecture

Similarly, SHG imaging of chronic inflammation in IL-10 mice reveals deterioration of collagen structure supporting the crypt wall, with crypt boundaries defined by a lower intensity signal than its corresponding healthy state (**Fig. 2A**), and the presence of disoriented fibrillar structure between crypts (**Fig. S2A**). Crypt wall thickness, as well as their diameters, increases and collagen ultrastructure is heterogeneous compared to the healthy IL-10 mice, from a diameter of 60µm to 85.3µm and from a wall thickness of 20.1µm to 37.8µm (**Fig. 2C** and **Fig. S2C-D**). Further inspection of the ECM structure using SEM reinforced this finding: naïve IL-10 crypt wall ECM appears fibrous, whereas in chronically ill IL-10 mice, the crypt wall ECM appears disintegrated and perforated, with fibrillary structures of different morphology (**Fig. 2B** and **Fig. S2B**).

#### Pre-symptomatic colitis ECM architecture

Remarkably, at the pre-symptomatic state of the DSS model (day 4), thickening or deterioration of some crypt walls occurs, with changes in crypt diameter, along with the appearance of fibrillar structures in the spaces between crypts (**Fig. 2A** and **2C** and pointed out by arrows in **Fig. S2A**), indicating remodeling events of the colonic ECM. Even more prominent ECM remodeling can be seen at this stage, using high-resolution SEM, as exposed fibrillar collagen on the inner crypt walls (**Fig. 2B**), due to destruction of the basement membrane layer coating the interstitial ECM (**Fig. S2B**).

Most surprising was the comparison of the healthy IL-10 mice to their WT counterpart and to DSS day 4. We observe a strong resemblance in ECM structure at high-resolution between the colon of IL-10 mice and that of the pre-symptomatic, day 4 DSS model, with the characteristic exposure of fibrils on the crypt wall and disrupted basement membrane, as pointed out by arrows in **Fig. 2B**, and shown in more detail in **Fig. S2B**. To test whether this observation is phenomenologically robust, we quantified the frequency of fibril exposure in the two pre-symptomatic states (day 4 of DSS and healthy IL-10), as a percentage of crypts that have exposed walls, by comparing to the frequency in healthy WT mice. We found that this structural change occurs in both models at a statistically significant rate, of ∼90% for day 4 and ∼75% for healthy IL-10 (**Fig. 2D**) compared to healthy WT (∼6%). Thus, IL-10 mice, which are prone to developing chronic inflammation, develop a “leaky” basement membrane as a quasi-stable pre-symptomatic state, compared to the dense mesh covering the WT murine colon (**Fig. 2B** and **Fig. S2B**).

Taken together, the structural similarity of the ECM between the two murine pre-symptomatic states, leads us to hypothesize that exposed fibrillar collagen due to the deterioration of the basement membrane, is one of the features of what we term the “pre-symptomatic ECM”.

### Disease onset is associated with a gradual loss of matrix rigidity

Having defined the structural and morphological differences of the matrix in pre-symptomatic states, we applied a second dimension of our analysis, which interrogates the changes in the mechanical properties of the matrix. To do this, we used mesoscale atomic-force microscopy (AFM) on decellularized ECM scaffolds derived from colons of mice from both experimental models, using pre-symptomatic (day 4 of DSS and healthy IL-10) and pathological (day 10 of DSS and chronically ill IL-10) states, in addition to healthy WT.

We used micron-sized probes to apply pressure to a 5mmX5mm (hydrated size) ECM scaffold. This permitted acquisition of the scaffolds’ mesoscopically averaged, statistically robust Young’s Modulus (YM) values[21, 22], which describe the collective contributions of nanoscale molecular components organized in micrometer-sized (70µmX70µm) domains in the ECM (see Methods for more details). Distributions of the measured YM values of colonic ECM samples are broad (**Fig. S3**), demonstrating the diversity of elastic properties within the same sample, and suggesting variability within the complex structure and composition of the ECM of each individual. Each mode in the distribution (**Fig. S3** and **Table S2**) represents the elastic properties of a region of the ECM with lateral dimensions (and thickness) of several microns, which we term “mechanical domains”. The median values and average median values of the YM of these mechanical domains are reported in **Fig. 2E**.

The rigidity of the ECM in healthy mice, both WT mice and IL-10, is similar and relatively high, with median YM values of about 11kPa. As inflammation is taking place, the colonic ECMs are generally softer than those of healthy colons, as indicated by the median YM (**Fig. 2E** and **Table S1**). Importantly, the structurally modified ECM in the pre-symptomatic state (day 4) of the DSS model has mechanical domains with an intermediate rigidity (YM=9kPa), with values between those of healthy (YM=10.5kPa) and acutely inflamed (YM=6kPa) states of DSS. The ECM of the chronically ill IL-10 mice (YM=3kPa) is much softer than that of the healthy IL-10 mice (YM=11kPa) (**Fig. 2E** and **Table S1**). Remarkably, chronically ill IL-10 mice possess a YM value that is approximately two-fold lower than that of the acutely inflamed colon (**Fig. 2E** and **Table S1**).

Altogether, our two-pronged analysis highlights a gradual softening of the ECM over the course of acute colitis development, peaking along with acute symptoms, which are accompanied by massive structural ECM damage.

### Matrisome protein compositional dynamics reflects the state of the colon

To understand the molecular outcome of the remodeling events that are evidently taking place in the ECMs of the two distinct pre-symptomatic states, we analyzed their protein composition – the *matrisome*[23]. We used mass-spectrometry (MS)-based analysis of the tissue proteome, followed by statistical analysis and computational modeling. Eight to ten colon samples were analyzed from each of the five states: healthy WT, the two pre-symptomatic states (i.e., day 4 DSS, healthy IL-10), and two types of intestinal inflammation (acutely ill day 10 of DSS, chronically ill IL-10). Out of 2045 total proteins identified in all states, 110 ECM and ECM-related proteins, comprising 5% of all identified proteins, were retrieved. Annotation as an ECM protein was made according to the Matrisome, an ECM protein cohort defined by Naba *et al* [23].

We identified the ECM proteins that are significantly changed in abundance among states (see Methods for details). A scheme summarizing our results is shown in **Fig. 3A**, and in more detail in **Table S3**. We found that the ECM composition of the healthy WT state is very different from that of both states of illness. We also detected a statistically significant proteomic compositional shift in both pre-symptomatic states compared to the healthy WT. A large portion of this protein abundance shift between WT and pre-symptomatic states (i.e., increase in Collagen alpha-2(I) chain, Fibrillin-1 and Transglutaminase-3; decrease in Mucin-2 and Lectin mannose-binding-1) is shared between the two pre-symptomatic states (**Fig. 3B**). Remarkably, these two states exhibit the highest compositional similarity of any pair, with no differentially abundant ECM proteins at all (**Fig. 3A**). Moreover, the ECM matrisomic shift from each pre-symptomatic state to its corresponding ill state also shows a shared signature, which includes the reduction of several structural components (e.g., collagens, Laminin and Fibulin-5) (**Fig. 3C**), corresponding to the structural damage observed in ill states (**Fig. 2**). The two ill states, though clinically and functionally distinct, are also quite similar in terms of differentially abundant proteins, as summarized in **Fig. 3A**.

**Figure 3.**
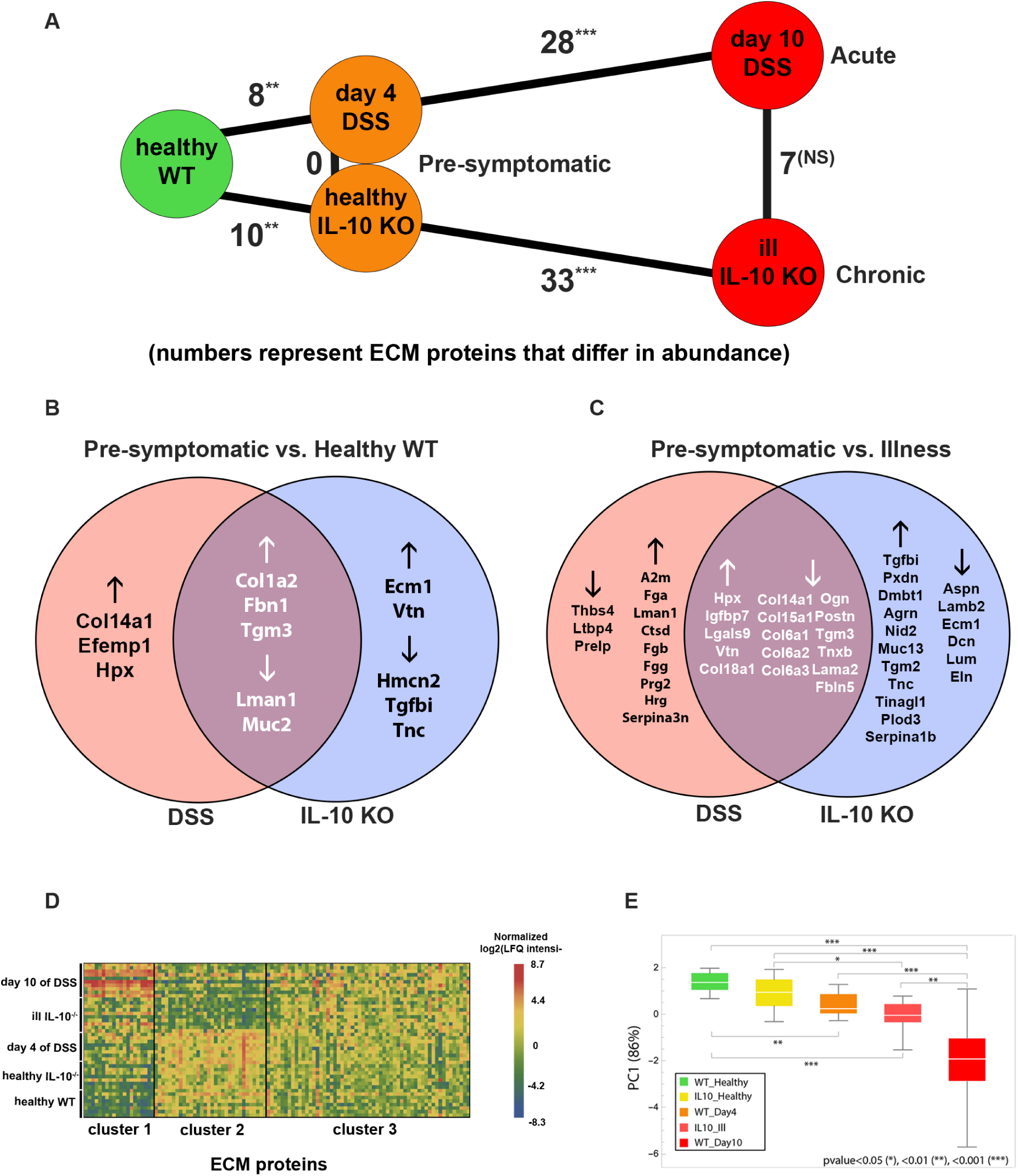
Pre-symptomatic states converge to a unique proteomic signature. (A) Scheme showing the number of differentially abundant ECM proteins, between pairs in the five different states, represented by lines stretched between two nodes of two states. Note that the two pre-symptomatic states are similar in terms of ECM composition. Statistical significance of amount of ECM proteins changing among all ECM proteins and all differentially abundant proteins, according to the sum of hypergeometric probabilities for achieving each number or above it, is indicated next to each comparison by asterisks. **P<0.01, *** P<0.001, NS=not significant. (B+C) Venn diagrams presenting the differentially abundant ECM proteins in the colon comparing healthy WT vs. the pre-symptomatic day 4 of the DSS model (pink) and healthy WT vs. pre-symptomatic healthy IL-10^-/-^ (blue) in B; and comparing pre-symptomatic day 4 of the DSS model vs. acutely ill day 10 of the DSS model (pink) and pre-symptomatic healthy IL-10^-/-^ vs. chronically ill IL-10-/-(blue) in C. The direction of change in abundance in pre-symptomatic states is indicated by arrows. Note the differentially abundant proteins shared among the two comparisons (purple) in both diagrams. The intersection between the differentially abundant proteins in both comparisons is statistically significant (p=1.0×10-4 in B and p=1.2×10-5 in C). Statistical significance in this figure was calculated by the sum of hypergeometric probabilities for achieving each number or above it. (D) MS data of 110(ECM proteins) x 44(samples at five clinical states) was clustered according to similar protein abundance by computing the squared Euclidean distance. Following, the data was partitioned into three clusters. Grouping according to clinical state is indicated on the x axis. Protein abundances were normalized to the mean of each protein. (E) PCA on the 3(clusters) x 44(samples) data after taking the mean of each cluster. The first PC explains more than 86% of the variance in the data. PC1 mostly (∼75%) consists of cluster 1. The PC1 values for each state are presented in a box plot. *P<0.05, **P<0.01, ***P<0.001. Note that the results show that ECM composition of clinical states can be projected onto a healthy-acutely ill axis, while highlighting the existence of the pre-symptomatic state.

To bridge the gap between omics data and tissue morphology and function, we further analyzed the data holistically, using unsupervised learning tools. These tools provide an unbiased analysis of the data, based on the variance of the input data, and not on the assigned identity, or clinical state of each sample. We identified three clusters of proteins with similar abundance patterns (**Fig. 3D**). We then used principle component analysis (PCA) on the means of each cluster in order to rank the different states according to their protein composition. Our analysis led to the formation of PC1, which is dominated (75%) by protease inhibitors (e.g., serpins) and coagulation factors (e.g., subunits of the fibrinogen complex), while the rest is composed from ECM structural proteins (e.g., collagens, proteoglycans and glycoproteins) (**Table S4**). PC1 reflects the tissue damage and ECM destruction characteristic of disease states. Strikingly, PC1 ranked the samples from the healthy WT to pre-symptomatic states (first healthy IL-10 and then day 4 DSS), followed by chronically ill IL-10 and finally, the acutely ill, DSS day 10 state. Thus differentiation of samples mostly according to the abundance of proteins in cluster 1 is sufficient to correctly distinguish between the five different states (**Fig. 3E)**. The two states at the edges of this ECM ranking, or hierarchy, represent the two extremes of their clinical states. We note that healthy IL-10 (as a chronic pre-symptomatic state) is closer to the “healthy edge” than the transient pre-symptomatic state of day 4 of DSS. This observation is consistent with the ECM rigidity analysis, described above.

In summary, the compositional analyses of all experimental samples provides a molecular signature for the “pre-symptomatic ECM” in both acute and chronically ill models, which is compositionally distinct from healthy WT or ill states, but is similar between inflammation models. Furthermore the compositional healthy-ill ranking arising from our unbiased computational analysis corresponds to the degree of structural damage that we observed via imaging while independently highlighting the existence of the pre-symptomatic state.

### Human IBD recapitulates murine matrisomic dynamics and validates biomarkers

In order to assess the clinical relevance of the identified biomarkers, we first tested whether the murine ECM proteomic analysis could be confirmed using an independent human dataset. One existing proteomic dataset comparing healthy donor biopsies with those of UC patients, deposited in the ProteomeXchange, was used for this experiment[24]. We examined the relative abundance levels of the proteins that were significantly differentially expressed between the healthy mice (both WT and IL-10) and their corresponding fully diseased sample types (**Fig. 4A**). A significant proportion of these proteins exhibit the same trends in relative abundance in the human dataset: proteins that are more abundant in ill mice than healthy mice are also more abundant in UC patients than in healthy donors, and vice versa (**Fig. 4B**).

**Figure 4.**
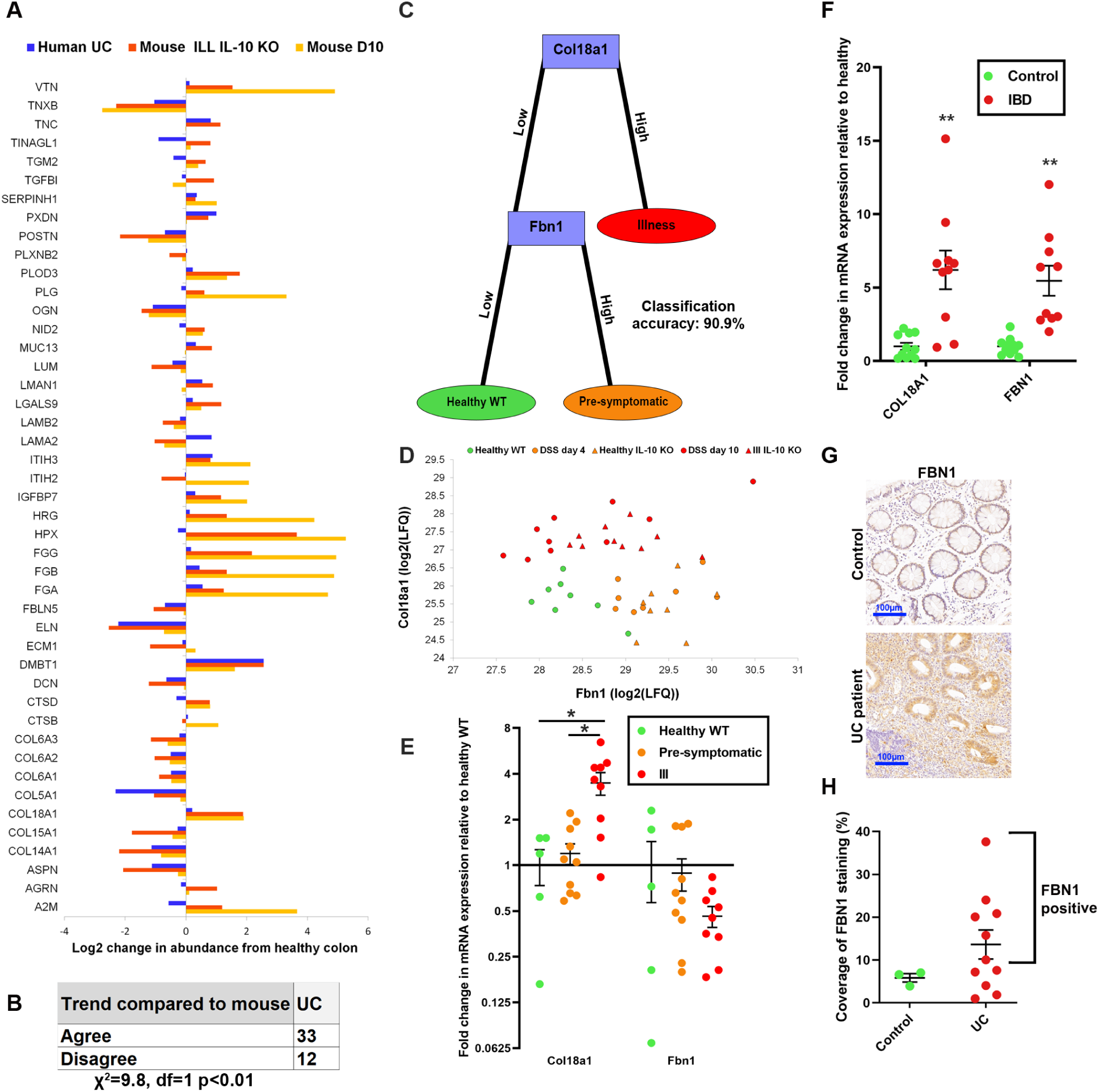
Identification of potential ECM biomarkers via machine learning and human validation. (A) Significantly different ECM proteins from MS analysis display similar trends in human dataset. Unpublished data from a proteomic data set of colon biopsies, Bennike *et al.*[24] were analyzed for changes in ECM protein abundance, comparing UC[24] patients to healthy controls. Proteins that changed significantly between healthy (WT or IL-10^-/-^) and ill (day 10 of DSS or chronically ill IL-10^-/-^, respectively) mice were chosen. (B) The directions of the observed changes in human were compared to that of the significant changes in mouse. Proteins that change in the same direction in both species were counted as those that “Agree” with the mouse data, compared to those that “Disagree”. A chi-square test of goodness-of-fit was performed to determine whether the rate of agreement in the direction of change was likely to occur by chance by comparing to a 50% (22.5) “Agree”-50% (22.5) “Disagree” distribution. (C) A method for identifying potential biomarkers by applying a classification algorithm. Samples were divided into Healthy WT, Pre-symptomatic (day 4 of DSS and healthy IL-10-/-) and Ill (day 10 DSS and ill IL-10-/-), and a J48 decision-tree classification algorithm [33] was applied to the data, along with a stratified 10-fold cross-validation. The two proteins chosen for the tree nodes (Col18a1 and Fbn1) are those that their abundance separate the three groups with the highest accuracy (90.9%) of all proteins. Col18a1 threshold: 26.67 log2(LFQ intensity). Fbn1 threshold: 28.76 log2(LFQ intensity). (D) Plot of all samples according only to the abundances of Col18a1 and Fbn1, demonstrating that the levels of the two proteins are good indicators of the tissue state. (E) Gene expression analysis of Collagen XVIII (Col18a1) and Fibrillin-1 (Fbn1) on independent mouse tissue samples of all three groups. Note, that Col18a1 transcription is significantly elevated in ill samples, while no significant changes were observed in Fbn1. n(Healthy WT)=5, n(Pre-symptomatic)=10, n(Ill)=9 (one outlier removed); *P<0.0167, t-test with Bonferroni correction. (F) Validation of biomarker transcript levels on an independent set of human samples. Biopsies and surgical resections of healthy donors or of inflamed mucosa of IBD patients were analyzed for mRNA expression levels of COL18A1 and FBN1. Both predicted markers indicated by the classification model performed on the murine models display significantly elevated levels of 6.2 fold (COL18A1) and 5.5 fold (FBN1) in inflamed IBD tissue compared to tissue from healthy donors (n(IBD)=10, n(Healthy)=11; *P<0.05, **P<0.01). (G) FBN1 IHC staining in rectal biopsies of healthy donors vs. inflamed regions in UC patients. Controls do not demonstrate FBN1 staining (FBN1 negative), while a subset (n=5) of UC patients display a strong epithelial staining (FBN1 positive). (H) Quantification of the area covered by FBN1 staining. The distribution of the FBN1 coverage in UC patients is broad, and half of the patients analyzed are FBN1 positive (above 10% coverage), such as the one presented in G.

In order to search for distinguishing features that can serve as potential biomarkers in our ECM proteomics data, we have utilized a machine-learning classification algorithm for analyzing our dataset. Identification of such distinguishing features would depend upon correct automated classification of samples into either “healthy WT”, “pre-symptomatic” or “ill” classes. The resulting model is illustrated in **Fig. 4C** (see legend for more details). The algorithm “chose” two proteins, Collagen XVIII/Endostatin (murine gene name: Col18a1) and Fibrillin-1 (murine gene name: Fbn1), as the nodes of the tree. Collagen XVIII is a ubiquitous non-fibrillar collagen, and a crucial, structurally complex basement membrane proteoglycan[25, 26]. The glycoprotein Fibrillin-1, is part of the microfibril-forming Fibrillin family that is important for connective tissue integrity and function[27]. Plotting all samples according to the abundances of these two proteins resulted in a clear separation between the healthy, pre-symptomatic and ill states (**Fig. 4D**). Of note, the pre-symptomatic state is defined by low levels of Col18a1 and high levels of Fbn1. Real-time qPCR analysis corroborated our findings regarding Col18 while no significant change in Fbn1 expression could be observed suggesting that upregulation of Col18a1 occurs at the transcriptional level, while Fbn1 is most likely regulated at the protein level (**Fig. 4E**). Remarkably, these observations of protein and transcript abundance exhibit similar patterns of relative abundance in clinically isolated human biopsies, as described below.

Next, we examined expression of these same putative biomarkers in a new set of human biopsies. For this, we compared mRNA expression levels of the human homologs, COL18A1 and FBN1, between healthy donor biopsies and tissues samples from IBD patients that were endoscopically and histologically identified as inflamed (sample list in **Table S6**). We found that both COL18A1 and FBN1 are significantly upregulated, in concordance with the decision tree applied to the proteomic data from our murine colitis samples (**Fig. 4F**). To corroborate this finding on the protein level and tissue localization, we also stained for Fibrillin-1 in inflamed rectal biopsies of pediatric UC patients with distal colitis. We have found that a subset (∼50%) of these patients display a strong epithelial staining of Fibrillin-1 (“FBN1 positive”), while the rest look like the control with little to no staining (**Fig. 4G-H**).

Taken together, analysis of published human proteomic datasets, qPCR and immunostaining of clinical biopsies, validate the clinical potential of the biomarkers Collagen XVIII and Fibrillin-1, which we identified using a classifier algorithm.

### Pre-symptomatic ECM changes coincide with epithelial gelatinase activity and myeloid cells bearing remodeling enzymes

Having observed that the most prominent structural ECM alteration, along with ECM softening, occurring at pre-symptomatic states involves perforation, and hence – degradation of the basement membrane underlying the epithelial cells of the colonic crypts and covering collagen fibrils, we set to examine the levels and cellular sources of mostly ECM proteases and gelatinolytic activity in these tissues. First, we tested mRNA expression in whole tissue specimens of both gelatinases -A and -B (MMP-2 and MMP-9), which primarily degrade basement membranes[28, 29]. We also tested other MMPs, LOX (a collagen crosslinking enzyme) and the tissue inhibitor of MMPs (TIMP)-1. We found that both MMP-2 and −9 were predominantly upregulated both at day 4 of the DSS model and in healthy IL-10 mice compared to healthy WT, and that the expression of TIMP-1, which inhibits MMP-2 and −9, was unchanged (**Fig. 5A**). Since the final regulatory step of MMPs that determines their net contribution to ECM remodeling[30] occurs outside of the cell, with their activation, we used the activity-based assay, *in situ* zymography[31] to localize active gelatinases in the tissue. Our analysis reveals that elevated (∼1.5-2 fold) gelatinolytic activity is detected in the crypts of the pre-symptomatic tissues, compared to healthy WT (**Fig. 5B-C**), and appears to localize to the epithelium.

**Figure 5.**
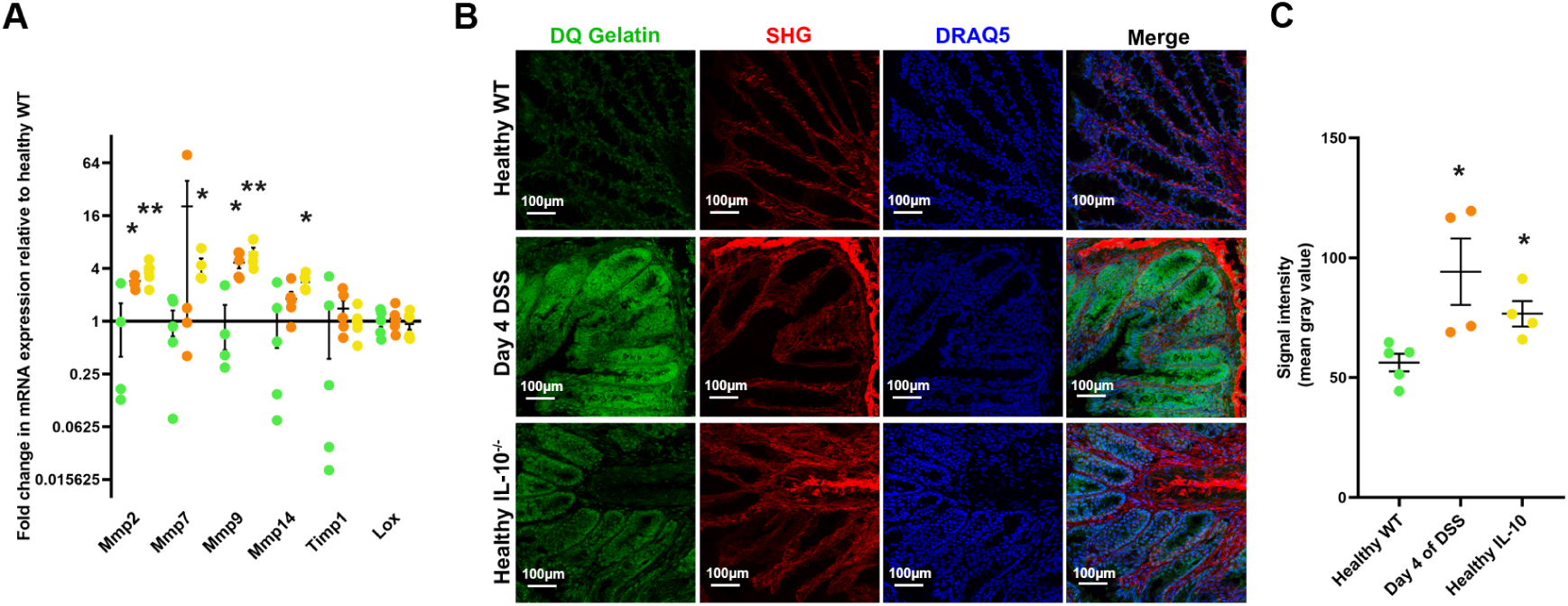
Pre-symptomatic tissues display elevated gelatinase expression and activity (A) Gene expression analysis of selected ECM remodeling enzymes (Mmp-2/-7/-9/-14) and inhibitor (TIMP1) at pre-symptomatic states. An independent set of mouse tissue samples of Healthy WT (green), Day 4 of DSS (orange) and Healthy IL-10^-/-^ (yellow) were used. Note, that gelatinases (Mmp-2/-9) are significantly upregulated at pre-symptomatic states. n=5 animals per state; *P<0.025, **P<0.005, t-test with Bonferroni correction. (B) In situ zymography with DQ Gelatin^TM^ on colon cross-sections, depicting elevated gelatinase activity (green) in the mucosa (possibly the epithelium) of the pre-symptomatic states, along with DRQ5-stained cell nuclei (blue) and SHG signal of fibrillary collagen (red). (C) Quantification of signal intensity in the mucosa, revealing that the elevation in gelatinase activity is statistically significant.

In order to investigate the cellular sources of ECM remodeling enzymes at the transient day 4 state, we used mass-cytometry analysis that allows us to sort cells according to a large number of markers. This yields a comprehensive picture of the relative frequencies of different cell populations that express proteins of interest. The results reveal that despite showing no signs of inflammation by standard clinical tools[17], and not having a significant change in the total proportion of immune cells in the tissue (**Fig. S4A**), mice on day 4 display a statistically significant elevation (0.38% of live cells) in the percentage of neutrophils in the colon compared to healthy WT (0.021% of live cells), as well as an observable, yet not fully pronounced, rise in monocyte levels (**Fig. 6A** and **Fig. S4B**). This finding demonstrates the earliest step of immune cell infiltration. Neutrophils and monocytes, carry a variety of remodeling enzymes, including LOX, MMP-7,-9 and −13 (**Fig. 6B-C**), thus indicating their contribution to the formation of the pre-symptomatic ECM, and the observed proteolytic activity in **Fig. 5B**. The data also reveals that MMP-2-expressing epithelial cells account for ∼10% of all living cells, though this proportion remains constant through day 4 (**Fig. S4F**), their contribution of MMP-2 may be augmented via expression levels per cell or activation levels (rather than the number of cells expressing it). These findings also suggest that MMP-9, an established marker of active IBD[6, 32], displays elevated activity at pre-clinical states as well, along with MMP-2. In addition, we found that neutrophils and monocytes secrete LOX, a collagen crosslinking enzyme, which may explain the thickening of the crypts walls at the pre-symptomatic state. Together with the multifaceted analysis of the previous sections, we show that elevated ECM enzymatic degradation from various cellular sources takes part in the structural ECM alterations that we observe at pre-symptomatic states.

**Figure 6.**
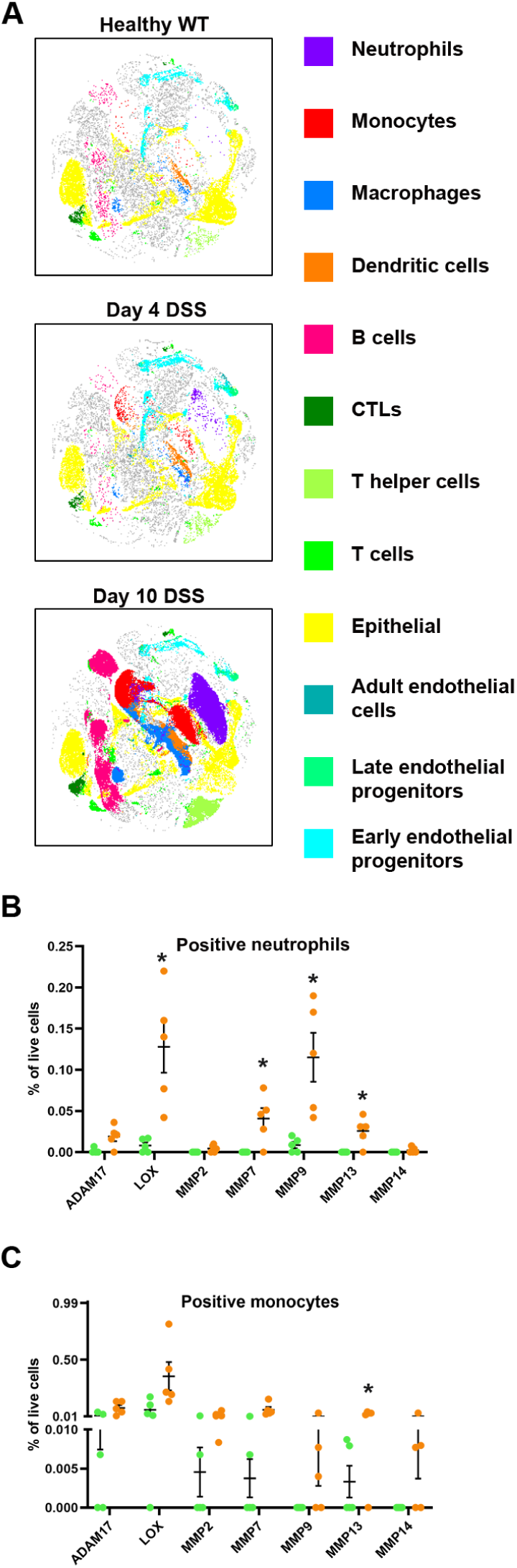
Enzyme-bearing immune cells participate in pre-symptomatic ECM remodelling. (A) tSNE analysis of mass cytometry demonstrating the changes in cell composition at each time point of the DSS acute colitis model. Each plot represents a pool of all five animals analyzed per state and each point corresponds to one cell. (B+C) Scatter dot plots based on the mass cytometry analysis depicting the changes in the relative frequencies (out of all live cells) of neutrophil and monocyte populations positive for the indicated remodeling enzymes at each state: healthy WT (green), day 4 of DSS (orange). Additional data (including neutrophils on day 10) from this analysis appears in **Fig. S4**. Note that a significant amount of infiltrating neutrophils carry ECM remodelers. Number of animals: n=5 per state; Bonferroni correction for three comparisons: *P<0.0167, **P<0.00333

## Discussion

The “pre-symptomatic ECM”, which we have identified and characterized, on multiple levels of analysis, herein, is a reproducible signature that is virtually identical in acute and chronic colitis models, and crucially, is distinct from that of the healthy WT tissue. Our analysis exposed several aspects and processes of remodeling, which potentially contribute to setting the stage for inflammation and understanding the nature of the challenge it imposes on the tissue.

Specifically, we identify the existence of a silent inflammatory tissue state by combining structural, mechanical and molecular system-level ECM analyses. We reveal that significant ECM remodeling precedes the appearance of detectible clinical symptoms, which is characterized by structural damage and perforation of the basement membrane, and changes in stiffness and proteomic composition. This remodeling is accomplished both by elevated production and activity of remodeling enzymes in the epithelium in both colitis models, along with sub-clinical immune-cell infiltration, in the acute model, which cannot be detected by conventional histopathological analysis. In addition, a recent report indicates that this pre-symptomatic ECM, in the specific case of the DSS colitis model, can be achieved via direct basement membrane damage inflicted by DSS[33], supporting our observation that basement membrane damage is part of the injury process in this model. Our unsupervised analysis of ECM compositions also showed that by narrowing our proteomic results into principal components (**Fig. 3E**) we can distinguish the pre-symptomatic state from all other states in the colitis model, enhancing our view that this is indeed a separate entity. Finally, our preliminary human IBD characterization recapitulates our observation of murine matrisomic dynamics and validates the found biomarkers.

Another discrete feature of the pre-symptomatic state highlighted by our characterization arises from the supervised learning analysis on the matrisome, which suggests a convergence of the five states examined into 1) healthy WT, 2) pre-symptomatic, and 3) ill. We identified Collagen XVIII/Endostatin and Fibrillin-1 as features distinguishing these states – indicating either ill or pre-symptomatic states, respectively. Neither of these proteins has been extensively studied in the context of IBD previously. Recently, a single report showed that upregulation of the FBN1 gene expression at diagnosis of pediatric IBD predicts the fibrostenotic complication of the disease[34]. This suggests the subset of patients with FBN1-positive epithelial staining in our study as candidates for long-term clinical follow-up, to determine whether this molecular ECM feature may have similar prognostic impact. Thus, machine learning as part of our compositional analysis has already demonstrated its potential for identifying relevant clinical markers and disease prognostic indicators.

Though IBD pathophysiology involves major tissue-damaging processes, which ultimately lead to the complications of the diseases[35–38], little is known to date regarding tissue remodeling events involving ECM before clinical presentation in these diseases. **Scheme 1** suggests a mechanism that connects our observed pre-symptomatic signatures with the promotion of inflammation.

**Scheme 1.**
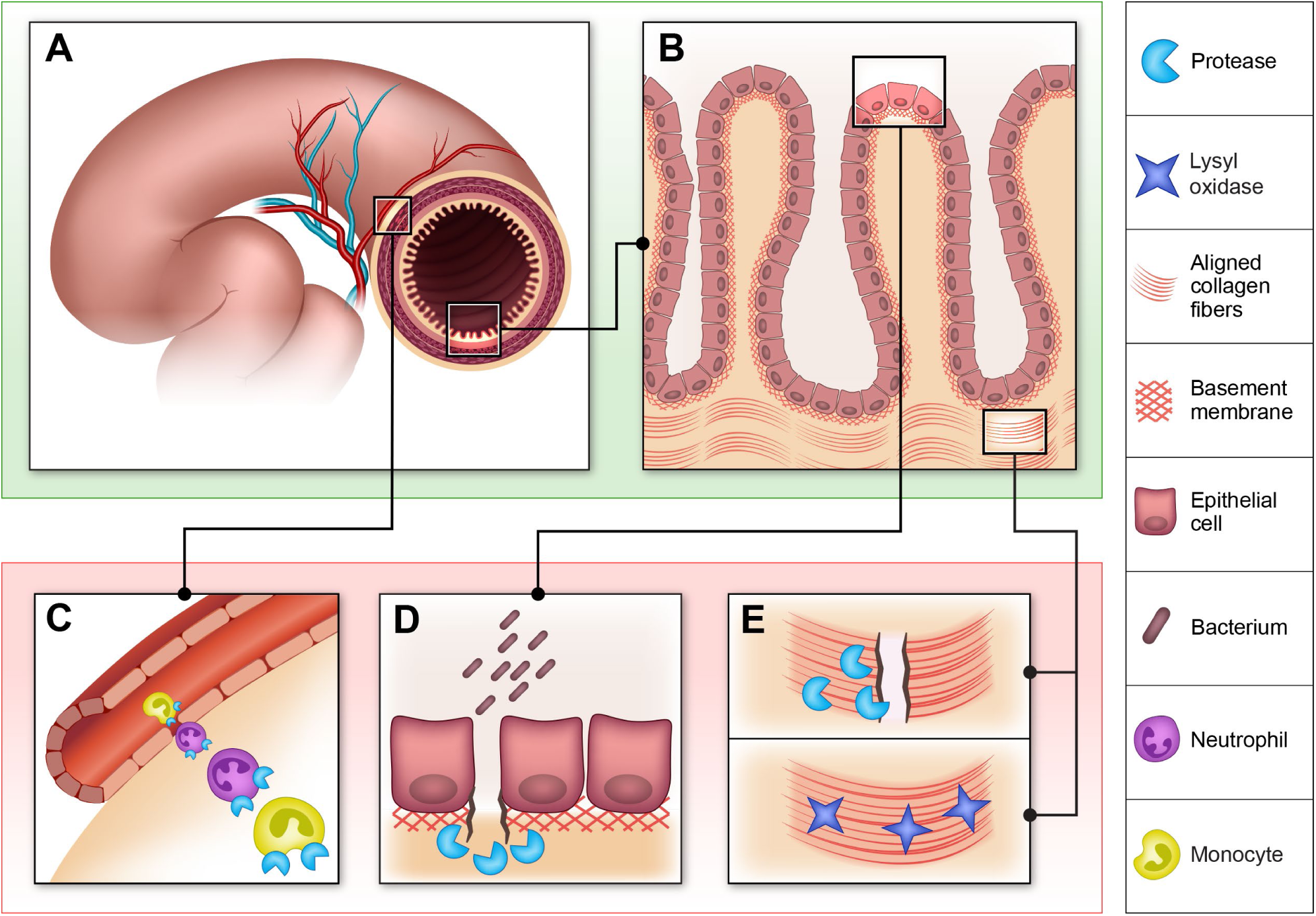
Pathophysiological contributions of the pre-symptomatic ECM. The scheme demonstrates how the ECM signatures found at the pre-symptomatic state lead to the inflammatory symptoms found in colitis. (A) Pre-symptomatic colon segment along with its vasculature. (B) Epithelial crypts composing the colonic mucosa. (C) Neutrophils and monocytes enhance immune cell infiltration capability via MMPs and LOX enzymes. (D) Protease activity, namely gelatinases in the epithelium, create a permeable mucosal barrier in the colon that grants gut microbes access to the internal environment. (E) Dual effect of increased MMP and LOX secretion along with increased fibrillary collagen secretion. This creates a deformed morphology characterized by both thick fibrous walls circumscribing the crypt in the presence of a perforated basement membrane.

Importantly, along with the convergence of pre-symptomatic states in two different colitis models, our findings are in line with current knowledge, stating that neutrophils bearing MMP-9 are often the first immune cell type to initiate the inflammatory process[39], and that the basement membrane is a key player in the barrier function of all mucosal tissues[40]. These strongly suggests that a similar pre-symptomatic ECM would exist in other inflammatory diseases, such as in the airway or other parts of the gastrointestinal tract. Therefore, we propose that the disruption to ECM integrity by a small number of pioneer neutrophils and monocytes, and the secretion of gelatinases by the epithelium, allows access to additional immune cell infiltration and bacterial antigens, which leads to full-blown inflammation.

In summary, our integrative approach permits a well-rounded characterization, encompassing multiple levels of detail, of the ECM – a complex and substantial material that comprises tissues. We conclude that by manifesting the pathophysiological state of a tissue, the ECM, in all of its complexity and properties, reveals silent disease-prone states that occur before clinical symptoms. The lessons from the observations we made in intestinal inflammation can be applied to other scenarios of inflammation due to the similarities of its onset in other mucosal tissues. We anticipate that this work will fuel mechanistic studies on the means by which early morphological and biomechanical ECM remodeling events leave the tissue vulnerable or prone to other pathologies such as cancer, fibrosis and inflammation in other organs. Identification of such additional silent pathological states by utilizing the ECM as a reporter will contribute to early diagnosis, managing relapse and intervention in tissue-damaging diseases.

## Methods

### Animals

Seven-week old C57BL/6 male mice were purchased from Envigo and were allowed to adapt for one week before experimental procedure. IL-10^-/-^ C57BL/6 mice from Jackson Laboratories were inbred at the Weizmann Institute of Science, and experiments were performed on six to eight-week old male mice. Healthy naïve IL-10^-/-^ were eight to twelve weeks old at time of harvest. All experiments and procedures were approved by the Weizmann Institute of Science animal care and use committees, protocol numbers 02230413-2, 06481012-1.

### Human samples

Biopsies were collected from healthy donors and from mucosa of IBD patients after providing informed consent in accordance with the ethical standards on Human Experimentation and the Declaration of Helsinki (protocol # 0725-14-TLV and # 3312-16-SMC). Samples were classified as being inflamed or normal by endoscopy and histology. Surgically resected inflamed colon samples from IBD patients were obtained from the Israel National Biobank (MIDGAM).

Samples were taken from both male and female subjects. The samples in each group for the RNA analysis are listed in **Table S6**. Samples used in RNA analysis were snap-frozen in liquid nitrogen until RNA extraction. Biopsies used for immunohistochemical stains were fixed in 4% paraformaldehyde.

### Intestinal inflammation induction and evaluation

Acute colonic inflammation was induced by administration of 1.25% DSS (MP Biomedicals LLC) in drinking water of C57BL/6 mice for seven days[15]. Chronic colonic inflammation was accelerated and synchronized by peroral administration of 200ppm piroxicam (Sigma-Aldrich ltd.) to IL-10^-/-^ C57BL/6 mice via supplementation in normal murine chow[16]. Mice were weighed 3 times a week over the course of the experiment. Colitis progression was evaluated over the course of the experiment using the Karl Stortz Coloview mini endoscope system and colonic inflammation was scored (0-15) as previously described[41]. Inflammation scores were categorized according to the following: healthy (0-4); mildly inflamed (5-7); inflamed (8-11); severely inflamed (12-15). Another form of inflammation evaluation was histological analysis by H&E staining of formalin-fixed paraffin-embedded tissues sections. The degree of histological damage and inflammation was graded in a blinded fashion by an expert pathologist. The following parameters were scored in the DSS-induced colitis model: amount of inflammation (0, none; 1, mild; 2, moderate; 3, severe; 4, accumulation of inflammatory cells in the gut lumen); percentage of colon area involved (0, none; 1, <=10%; 2, 10%-30%; 3, 30%-50%; 4, 50%-90%; 5, >90%); depth of inflammation and layers involved (0, none; 1, mucosa only; 2, mucosa and submucosa; 3, limited transmural involvement; 4, transmural); ulceration (0, <30%; +1, >30%; - 1, regeneration). The overall histological score was the sum of the three parameters (maximum score of 14). Histopathological scoring for the IL-10 model was performed in a similar fashion, but only considering the amount of inflammation and percentage of colon area involved (maximum score of 9). IL-10 mice lose weight only in early phases of colitis development, and the weight is regained as chronic colitis is established (**Fig. 1E**).

### Mass cytometry and cell sorting

Cell isolation of colon samples was performed as previously described[42]. In brief, colons were harvested on the day of the experiment from healthy WT mice, along with those of day 4 and day 10 time points of the DSS model, were carefully separated from the surrounding fat and lymphatic vessels and cleaned from feces with cold PBS. Then, 2cm colon pieces were minced and immersed into RPMI-1640 medium (GIBCO, 21875034) containing 10% FBS, 2 mM HEPES, 0.05mg DNase (Roche, 04716728001), and 0.5mg/mL of collagenase VIII (Sigma-Aldrich, C2139) and placed for 40 minutes at 250 rpm shaking at 37°C and mashed through a 70µm cell strainer.

Following, cells were stained according to a previously published protocol[43]. Individual mice cell suspensions were stained with Cell-ID Cisplatin 0.125 µM for viability and fixed using Maxpar® Fix I Buffer. Samples were then permeabilized using Maxpar® Barcode Perm Buffer and then barcoded using The Cell-ID™ 20-Plex Pd Barcoding Kit, allowing us to join samples for antigen staining. Antibodies used for staining are listed in **Table S7**. Before analyzing, the cell suspension was incubated with Cell-ID Interculator Iridium for 20 minutes.

Cells were analyzed with a cyTOF2® mass cytometer (Fluidigm). Results were normalized and debarcoded using Fluidigm cyTOF software[44]. CyTOF results gating and further analysis was done with FlowJo software (FlowJo, LLC) and cell populations were defined according to the markers listed in **Table S8**. tSNE analysis was done using the viSNE application in the Cytobank web platform. tSNE (t-Distributed Stochastic Neighbor Embedding) is a machine-learning algorithm used to cluster multivariate data into a 2D representation.

### Decellularization of colonic tissue

Decellularization was performed as previously described[45]. Samples were frozen in ddH_2_O and subjected to six freeze-thaw cycles with replacement of ddH_2_O each time. Subsequently, samples were immersed in 25mM NH_4_OH (Merck) for 20 minutes. Finally, isolated ECM scaffolds were washed six times in ddH_2_O and stored at 4°C until use.

### Second-harmonic generation (SHG) microscopy

Native snap-frozen murine colon samples were thawed in PBS and imaged using a two-photon microscope (2PM:Zeiss LSM 510 META NLO; equipped with a broadband Mai Tai-HP-femtosecond single box tunable Ti-sapphire oscillator, with automated broadband wavelength tuning 700–1,020 nm from Spectraphysics, for two-photon excitation). For second-harmonic imaging of collagen, a wavelength of 800-820 nm was used (detection at 390-450nm). Quantification of crypt diameter was done using ImageJ software (Research Service Branch, NIH).

### Scanning electron microscope (SEM)

Decellularized mouse colon tissues were used and prepared as described in [46]. Then, samples were subjected to critical point dehydration (CPD) and attached to a carbon sticker. Finally, samples were coated with gold palladium, and imaged under SEM Ultra (Carl Zeiss international).

### In situ zymography

In situ zymography was conducted as previously described[31] on unfixed 10μm transverse mouse colon sections using DQ Gelatin^TM^ (Invitrogen) from pig skin. Following, sections were fixed with 4% paraformaldehyde, and cell nuclei were stained with DRAQ5 (Thermo Scientific). Finally, mounting solution (Immu-Mount^TM^ Thermo Scientific) was added and slides were covered with a coverslip. The slides were imaged under a Leica TCS SP8 microscope (both confocal and multiphoton). DQ gelatin was excitated at 488nm and its emission was detected at 515nm.

### Testing elastic properties of ECM samples by Atomic Force Microscopy

The AFM analysis was carried out on rehydrated 5mmX5mm ECM samples derived from mouse colon tissues. Typically, three-four samples were analyzed for each inflammatory condition. The rehydration process was as follows: ECM samples were grossly dried and attached to glass coverslips (diameter 15 mm) by means of a thin bi-adhesive tape. Samples were then attached to the bottom of Petri dishes (Greiner Bio-One) and left overnight in an evacuated desiccator in order to dry out, so to improve spreading and adhesion of ECM on tape. Prior to AFM measurements, the Petri dish hosting the ECM sample was filled with ddH_2_O and the ECM was allowed to rehydrate for 30 minutes at room temperature. Measurements were carried out at room temperature.

To accomplish the nanomechanical characterization of the ECM samples, a Bioscope Catalyst AFM (Bruker) was used to collect series of force vs distance curves[47, 48], recording the local deformation of the sample induced by the AFM probe. According to a recently published protocol[22], we used monolithic borosilicate glass probes consisting in micrometer-sized spherical glass beads with radius R=9-10 µm attached to silicon cantilevers with a force constant k=0.25-0.35 N/m. The probes were produced and characterized as previously described[21].

Each set of force curves (a force volume) consisted of a 16×16 array of curves acquired on a 70μm × 70*μ*m area. Ten force volumes were typically recorded on each ECM on macroscopically separated regions. All measurements were performed with the following parameters: 4096 points per curve, ramp length *L=10 μm*, maximum applied force *F = 60-70nN*, and ramp frequency *f=1.10 Hz*. Typically, indentations up to 2-3 μm were obtained. Data processing of force volumes was carried out in Matlab according to the general protocol described previously[22]. The values of the Young’s Modulus (YM) were extracted by fitting the Hertz model to the experimental data[47, 48]. A first very soft indentation region (0-40% of total indentation) was excluded, in order to separate the possible contribution of loosely-bound superficial layers from those of the bulk ECM. Artifacts derived from an ill-defined contact area between sample and probe, like boundaries of colonic crypts or more generally crypts with characteristic dimensions comparable to, or larger than the probe diameter, were identified and discarded.

The distributions of YM values of the ECMs from mice turned out to be the envelope of several nearly lognormal modes, representing the major contributions to the overall ECM rigidity and originating from micro-scale domains that the AFM probe was able to resolve. Each mode in the distributions represents the elastic properties of a region of the ECM with lateral dimensions (and thickness) of several microns, which may define a structural and functional domain of the ECM. Multi-Gaussian fit in semilog10 scale allowed identification of the peak value E’ and the geometric standard deviation 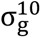 of each lognormal mode; from these values the median value E_med_ and the standard deviation of the median σ_med_ were evaluated for all modes as E_med_ = 10^E′^ and 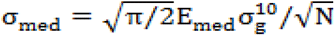 [49], N being the number of force curves in each mode. The effective rigidity of each ECM sample was characterized by the weighted average of median values 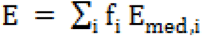, using the fraction *f*_i_ = *N*_i_/*N*_tot_ of force curves in the mode as the weight; the total error σ_E_ associated to E was calculated by summing in quadrature the propagated error of the medians 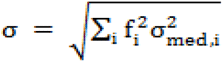 and an effective instrumental relative error σ_instr_ = 3 % : 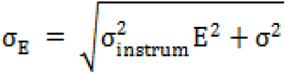. The average median values of the YM of the different states of inflammation have also been evaluated; the corresponding error has been calculated as the standard deviation of the mean summed in quadrature with the propagated σ_E_.

### ECM proteomics by liquid chromatography–tandem mass spectrometry (LC-MS/MS) analysis

Tissue slices from colons of WT or IL-10 mice at different stages of the inflammation models were immersed in a solution containing 50% trifluoroethanol (TFE), 25mM ammonium bicarbonate (Sigma Aldrich) and 15mM dithiothreitol (DTT) and coarsely homogenized by repeated cycles of boiling, freezing and sonication. Subsequently, samples were shaken at 30°C, 1400 rpm for 30 minutes, followed by the addition of iodoacetamide (Sigma Aldrich) to a final concentration of 25mM and further shaking for 30 minutes at 30°C and 1400 rpm. TFE was then diluted to 25% with 50mM ABC, LysC (Wako Chemicals, 1:100 lysC:protein ratio) and sequencing-grade modified trypsin (Promega, 1:50 trypsin:protein ratio) was added and incubated overnight at room temperature. On the following morning, more trypsin was added (1:80 trypsin: protein ratio) for 4 hours. Peptide mixtures were purified on C18 stage tips. Eluted peptides were loaded onto a 50 cm long EASY-spray reverse phase column and analyzed on an EASY-nLC-1000 HPLC system (Thermo Scientific) coupled to a Q-Exactive Plus MS (Thermo Scientific). Peptides were separated over 240 minutes with a gradient of 5−28 % buffer B (80% acetonitrile and 0.1% formic acid). One full MS scan was acquired at a resolution of 70,000 and each full scan was followed by the selection of 10 most intense ions (Data dependent Top 10 method) at a resolution of 17,500 for MS2 fragmentation.

Raw MS data was analyzed with MaxQuant software[50, 51] (version 1.5.2.18) with the built-in Andromeda search engine[52] by searching against the mouse reference proteome (UniprotKB, Nov2014). Enzyme specificity was set to trypsin cleavage after lysine and arginine and up to two miscleavages were allowed. The minimum peptide length was set to seven amino acids. Acetylation of protein N termini, deamidation of asparagine and glutamine, and oxidation of methionine were set as variable modifications. Carbamidomethylation of cysteine was set as a fixed modification. Protein identifications were sorted using a target-decoy approach at a false discovery rate (FDR) of 1% at the peptide and protein levels. Relative, label-free quantification of proteins was performed using the MaxLFQ algorithm integrated into MaxQuant environment with minimum ratio count of two[53].

### Computational analysis of MS data

Classification was done using a J48 decision tree algorithm[54] with a stratified 10-fold cross-validation in the Weka software for machine learning (University of Waikato). As input for this classifier, we used the ECM proteomic data (i.e., 110 matrisome proteins) described above, comprising 8 healthy WT samples, 16 pre-symptomatic samples (day 4 of DSS and healthy IL-10) and 20 ill samples (day 10 of DSS and ill IL-10), where each sample came from a different mouse. Clustering according to squared Euclidean distance and principal component analysis (PCA) were performed in Matematica (Wolfram Research).

### Quantitative Real Time PCR on mouse and human colon tissues

Colon tissues were homogenized using a bead beater homogenizer. Total RNA from was isolated using the PerfectPure RNA Tissue Kit (5 Prime GmbH) according to the manufacturer’s protocol. 0.6-1 μg of total RNA is reverse transcribed using High Capacity cDNA Kit (Applied Biosystems inc.). qRT-PCR was performed using SYBR Green PCR Master Mix (Applied Biosystems inc.) on an ABI 7300 instrument (Applied Biosystems). Values are normalized to an Actin-β (Actb) control in mouse tissues and to Hypoxanthine-guanine Phosphoribosyltransferase (HPRT) and Ubiquitin C (UBC) controls in the human tissues. Primer sequences are listed in **Table S5**. Data is presented as mean fold change compared to either healthy WT in mouse tissues or to normal, non-inflamed, human tissues using the 2^-ΔΔCT^ method[55]. Mice samples were grouped according to general state: (i) healthy WT, (ii) pre-symptomatic (day 4 of DSS and healthy IL-10) and (iii) ill (day 10 of DSS and ill IL-10). Standard error of the mean (s.e.m) was calculated on the 2^-ΔCT^ data, as was the statistical analysis.

### Immunohistochemical (IHC) stain for Fibrillin-1

Tissues were fixed in 4% paraformaldehyde and embedded in paraffin, and serial 5-μm sections were prepared from the whole biopsy for IHC. Sections were de-paraffinized and epitope retrieval was performed. Sections were incubated for 1 h with anti Fibrillin1 (Abcam), diluted 1:100, at RT followed by secondary antibody (HiDef Detection Polymer, 954D, Cell Marque, Rocklin, CA, USA). A peroxidase substrate kit (SK-4100, Vector Labs, USA) was used as a chromogen and hematoxylin as a counterstain. Tissue exposed only to the secondary antibody was used as negative control.

Staining coverage was quantified as the mean percentage of area covered by staining out of the total image in 3-10 fields of view per sample. The analysis was carried out using ImageJ software (Research Service Branch, NIH) by applying color deconvolution for DAB staining, followed by a consistent binary threshold.

### Quantification and Statistical Analysis

#### General

Statistical parameters including the exact value of n, the definition of center, dispersion and precision measures (mean ± s.e.m) and statistical significance are reported and portrayed in the figures and the figure legends. Wherever outlier removal is mentioned, outliers are considered values that are over 3 median absolute deviations distance from the median, which is a robust outlier detection method[56]. Data is judged to be statistically significant when P < 0.05 by a two-tailed student’s t test or with a Bonferroni correction of 0.05/m, where m=number of pairwise comparisons. In figures, asterisks denote statistical significance as calculated by student’s t test (∗, P < 0.05 or 0.05/m; ∗∗, P < 0.01 or 0.01/m; ∗∗∗, P < 0.001 or 0.001/m) in Microsoft Office Excel.

#### Statistical analysis of MS data

Bioinformatics analysis was performed with the Perseus program version 1.5.1.4[57]. The proteomic data was first filtered to remove the potential contaminants, proteins only identified by their modification site and reverse proteins. Next, the intensity values were log2 transformed and data was filtered to have at least three valid values in each group. Missing values were imputed based on normal distribution and differentially abundant proteins were chosen according to a student’s t-test with a Bonferroni correction for multiple (6) pairwise comparisons (corrected α=0.05/6=0.0083), and a 0.05 permutation-based false discovery rate (FDR) cutoff for analysis of multiple proteins[58]. Statistical significance of intersections in Venn diagrams and proportion of differentially abundant ECM proteins out of all differentially abundant proteins was assessed using a hypergeometric probability density function in Matlab (Matworks inc.).

## Acknowledgments

We thank Dr. Ori Brenner for assistance in histopathological scoring of intestinal inflammation. Also, we thank Dr. Miriam Rosenberg for helpful comments on the text and styling.

## Funding

IS is the Incumbent of the Maurizio Pontecorvo Professorial Chair and has received funding from the Israeli Science Foundation (1226/13), the European Research Council AdG (THZCALORIMETRY - DLV-695437) and the USA-Israel Binational Science Foundation (712506-01), the Ambach fund, and the Kimmelman center at the WIS. AP thanks the Dept. of Physics of the University of Milano for financial support under the project “Piano di Sviluppo dell’Ateneo per la Ricerca 2014”, actions: “A. Upgrade of instrumentation”, and “B. Support to young researchers”. AS and TG are funded by the Israel Ministry of Science Technology and Space.

## Author contributions

Conceptualization, ES and IS; Methodology, ES, RA, IA, AS, MA, LP and VG; Formal Analysis, ES, IA, AS, MA, LP and VG; Investigation, ES and RA; Resources, OM, NG, SF, LW, DSS, CV, AP, TG, PM; Writing – Original Draft, ES; Writing – Review & Editing, ES, IA, ISo, IS; Visualization, ES, MA, LP, VG and AP; Supervision, ISo, CV, DSS, AP, PT, TG, PM, CL, UA and IS; Funding Acquisition, IS, AP, TG, PM.

## Declaration of Interests

The authors declare no competing interests.

## Data availability

The MS proteomics data have been deposited to the ProteomeXchange Consortium via the PRIDE partner repository with the dataset identifier PXD004740 and can be viewed by logging in with the following details: Username; Password: hKx9zFSu.

## Supplemental Information

**Fig. S1.**
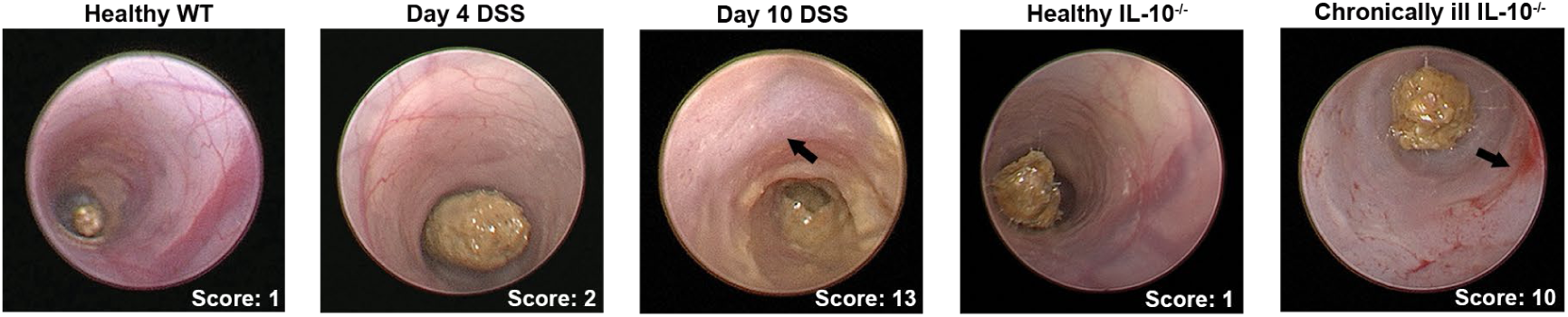
Endoscopic images. Live mice were imaged at the indicated states of the two models and inflammation scored as described in the Methods. The arrow in image of day 10 of the DSS model indicates the thick, granulated opaque mucosal surface. Chronically ill IL-10-/-mice predominantly display vascular disturbances and bleeding, as indicated by arrow.

**Fig. S2.**
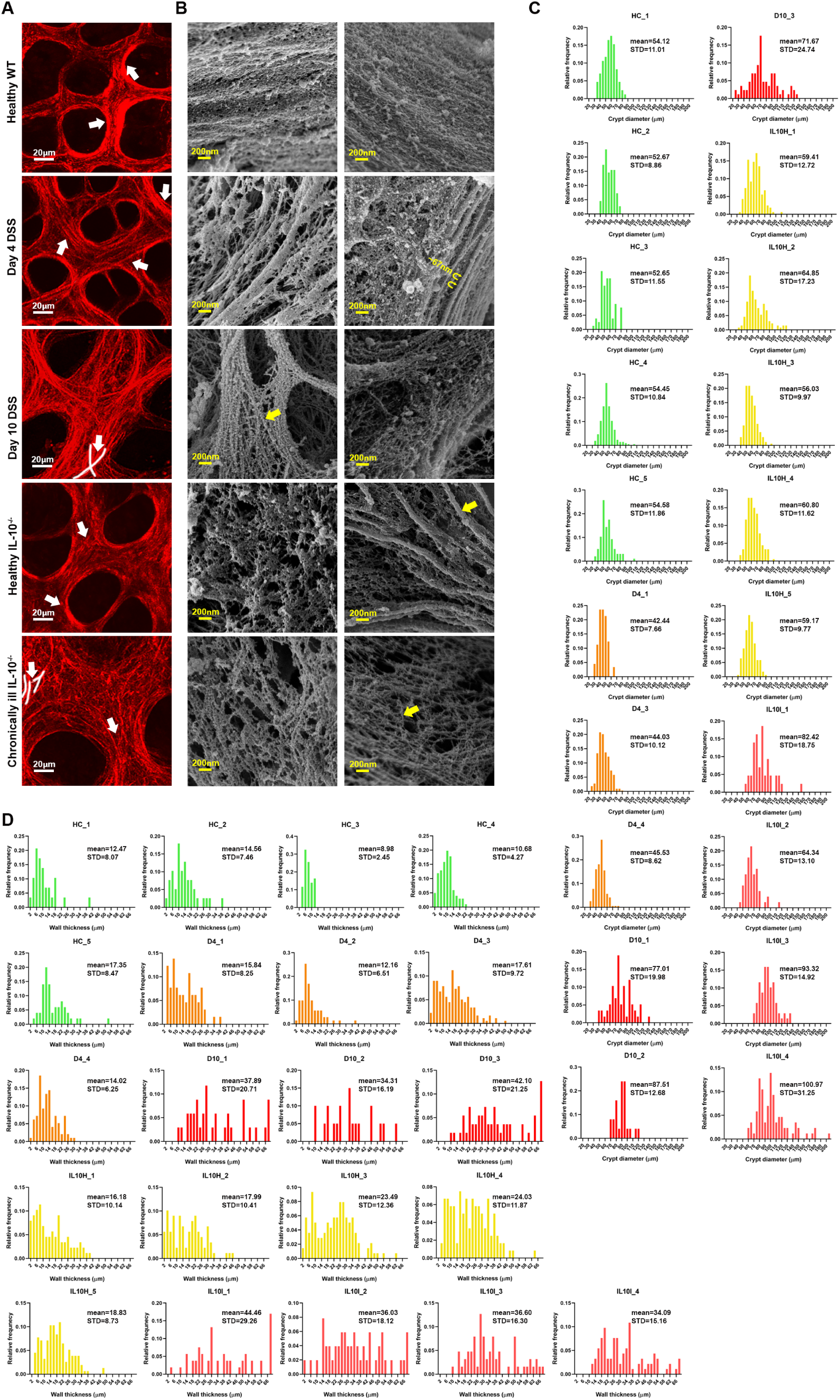
High-magnification images of SHG and SEM for emphasizing the structural changes and features of the ECM in all five tissue states. (A) Zoomed-in images of the ones in Fig. 2A. The higher magnification images allow us to demonstrate the changes in collagen fibers, which are pointed out by arrows. In healthy WT mice collagen is condensed around crypt borders, while fibers are somewhat splitting on day 4 of the DSS model as well as in healthy IL-10 mice. We see how the fibers become even more disoriented in both ill states, which is highlighted by white lines that follow selected intersecting fibers. This indicates that collagen fibers are splitting as a result of enzymatic degradation [59]. The high magnification also emphasizes the loose collagen structure characteristic of the chronic illness that contains thinner fibrils, which do not emit a strong SHG signal. In contrast, day 10 of the DSS model has fibers that emit a strong signal, indicating the coincidence of degradation (splitting) and deposition. (B) Additional SEM images, each image came from a different mouse. The images demonstrate the deterioration observed when comparing healthy WT to the two pre-symptomatic states (day 4 of DSS and healthy IL-10) in the ECM network that comprises the first layer that underlies the epithelium, which is most likely the basement membrane, since it resembles a collagen type IV network[20]. Not only are collagen fibrils exposed, as indicated by their characteristic 67nm D-banding (pointed out on day 4 in this panel), but also the network that coats them is not condensed and contains holes, both at day 4 and in healthy IL-10 mice. In addition, the changes in the observed fibrils as illness develops are pointed out by the arrows – while we see thick fibers at pre-symptomatic states, those are not present in acute (day 10) or chronic colitis (ill IL-10), and only thin, disoriented fibrils can be observed. (C+D) Histograms depicting the distributions of crypt diameters, in **C**, or wall thickness, in **D**, of each sample whose mean and STD can be found in the plots in Fig. 2C.

**Fig. S3.**
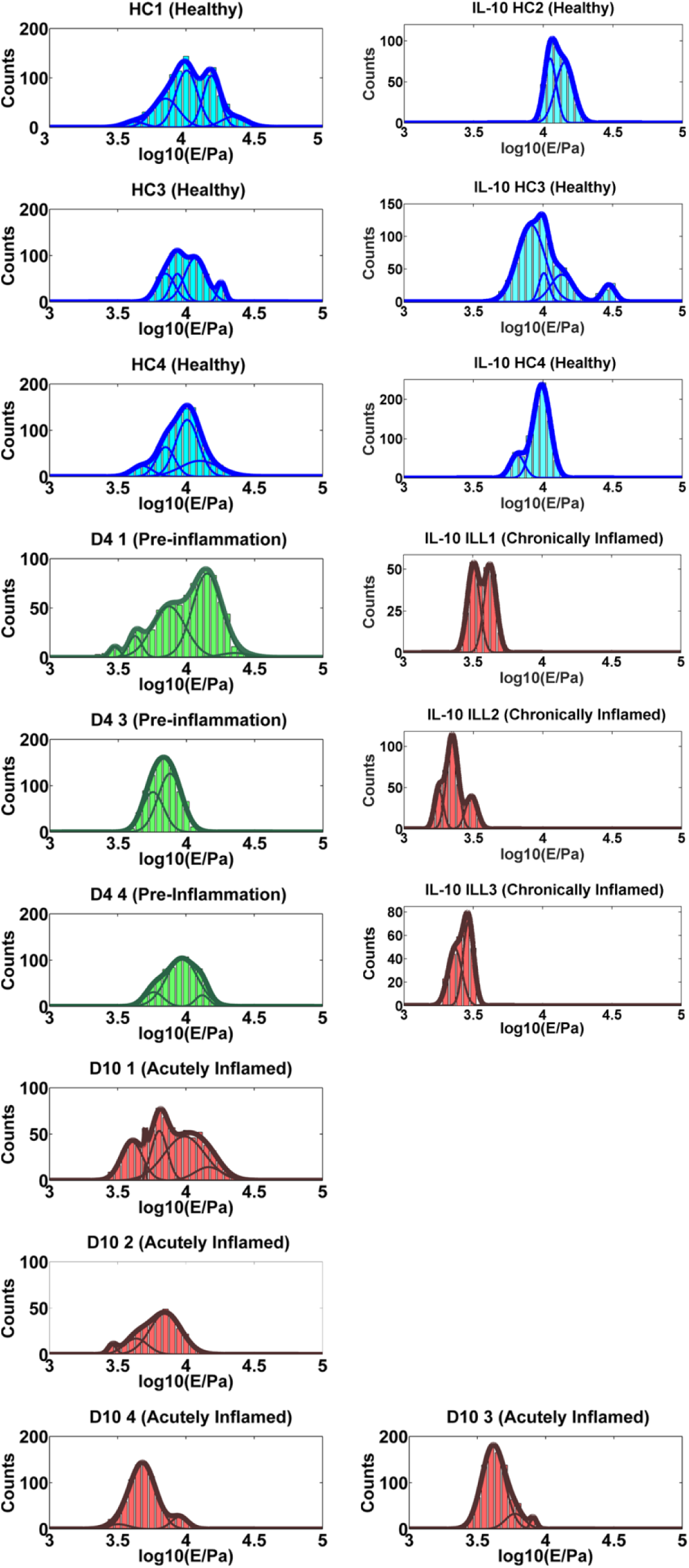
Collective distributions of Young’s Modulus values (E) measured on ECMs derived from colons of mice at three different states of the DSS-induced colitis model and two states of the PAC IL-10^-/-^ model (HC: healthy colon; D4: pre-symptomatic; D10: acutely inflamed; IL-10 HC: healthy naïve IL-10^-\-^; IL-10 ILL: chronically inflamed IL-10^-/-^). Each sample was derived from a different mouse. Thick lines represent the result of the superposition of the underlying major contributions, highlighted by thin continuous lines. Single contributions are detected by fitting the envelope of several Gaussian profiles to data in semilog10 scale. Under the hypothesis that the underlying distributions are log-normal, the peaks of the Gaussians represent the median values of the Young’s Modulus. Different contributions arise from the fact that each ECM is tested in different locations, therefore multiple modes witness the mechanical and structural heterogeneity of ECMs.

**Fig. S4.**
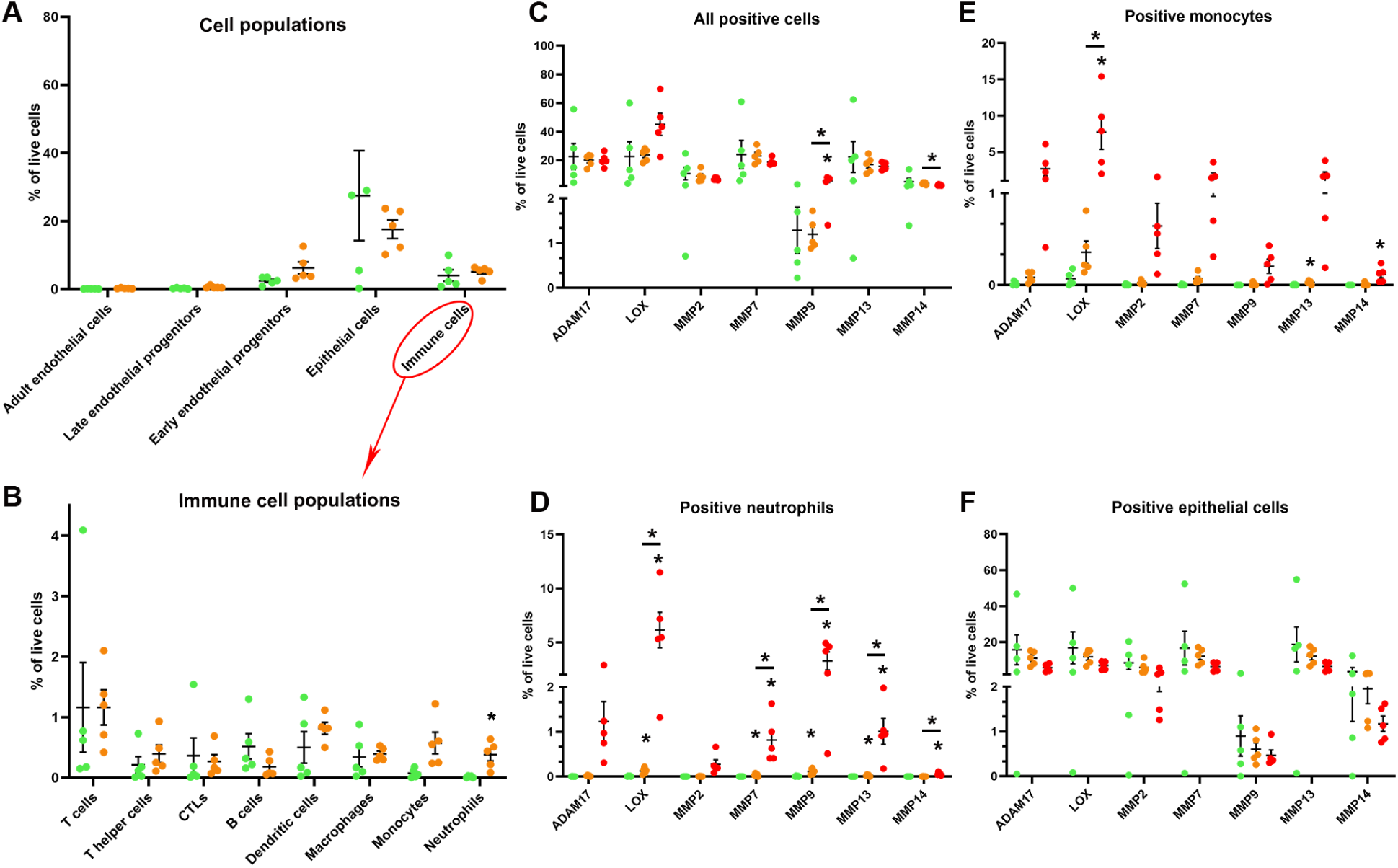
(A-B) Scatter dot plots based on the mass cytometry results depicting the changes in cell population relative frequencies (out of all live cells) comparing healthy WT (green) and day 4 of DSS (orange). Note that the only statistically significant change in cell composition on day 4 is a slight increase in the relative frequency of neutrophils. Number of animals: n=5 per state; Bonferroni correction for three comparisons: *P<0.0167, **P<0.00333. (C-F) Scatter dot plots based on the mass cytometry results depicting the changes in the relative frequencies (out of all live cells) of different cell populations positive for the indicated remodeling enzymes (C-all positive cells, D-neutrophils, E-monocytes and F-epithelial cells) at each state: healthy WT (green), day 4 of DSS (orange), day 10 of DSS (red). Number of animals: n=5 per state; Bonferroni correction for three comparisons: *P<0.0167, **P<0.00333.

**Table S1.** The statistical indicators and associated errors derived from the single-mode analysis for the collective distributions of Young’s Modulus values of ECMs represented in **Fig. S3**, which are plotted in **Fig. 2E**. P values when comparing average medians: Healthy WT vs. Day 10 (P<0.01); the differences between Healthy WT vs. Day 4 and Day 4 vs. Day 10 are not statistically significant (P=0.30 and P=0.09, respectively); Healthy IL-10^-/-^ vs. Ill IL-10^-/-^ (P<0.002). STD – standard deviation. All data are expressed in kPa units.

| Sample | | Young's Modulus / kPa<br>(weighted median ±<br>propagated errors) | Young's Modulus / kPa<br>(average median ±<br>effective STD of the mean) |
| --- | --- | --- | --- |
| Healthy WT (day 0) | HC 1 | 11.61 ± 0.25 | 10.55 ± 0.49 |
| | HC 3 | 10.39 ± 0.26 | |
| | HC 4 | 9.65 ± 0.25 | |
| Pre-Symptomatic (day 4) | D4 1 | 11.07 ± 0.39 | 9.03 ± 1.01 |
| | D4 3 | 6.86 ± 0.25 | |
| | D4 4 | 9.15 ± 0.36 | |
| Acutely inflamed (day 10) | D10 1 | 8.23 ± 0.24 | 6.04 ± 0.72 |
| | D10 2 | 6.31 ± 0.24 | |
| | D10 3 | 4.47 ± 0.15 | |
| | D10 4 | 5.14 ± 0.18 | |
| Healthy IL-10<sup>-/-</sup> | IL-10 HC2 | 12.94 ± 0.30 | 11.00 ± 0.90 |
| | IL-10 HC3 | 10.84 ± 0.20 | |
| | IL-10 HC4 | 9.21 ± 0.24 | |
| Chronically ill IL-10<sup>-/-</sup> | IL-10 ILL1 | 3.71 ± 0.06 | 2.89 ± 0.34 |
| | IL-10 ILL2 | 2.32 ± 0.03 | |
| | IL-10 ILL3 | 2.64 ± 0.04 | |

**Table S2.** Major contributions to the broad statistical distributions of the Young’s modulus values (see Methods section for details). STD – standard deviation. All data are expressed in kPa units.

| Sample | Young's Modulus / kPa<br>(median ± STD of the median) | | | | | |
| --- | --- | --- | --- | --- | --- | --- |
| Healthy WT 1 | | 4.17 ± 0.14 | 7.08 ± 0.12 | 10.12 ± 0.11 | 15.33 ± 0.15 | 22.40 ± 0.60 |
| Healthy WT 3 | | | 7.08 ± 0.19 | 8.63 ± 0.10 | 18.20 ± 0.21 | |
| Healthy WT 4 | | 4.76 ± 0.11 | 7.08 ± 0.09 | 10.20 ± 0.11 | | |
| | | | | 12.60 ± 0.33 | | |
| Day 4 1 | 2.98 ± 0.07 | 4.21 ± 0.08 | 7.42 ± 0.15 | | 14.16 ± 0.20 | 22.66 ± 1.46 |
| Day 4 3 | | | 5.68 ± 0.07 | | | |
| | | | 7.59 ± 0.08 | | | |
| Day 4 4 | | | 5.80 ± 0.11 | 9.35 ± 0.13 | 13.16 ± 0.24 | |
| Day 10 1 | | 4.02 ± 0.07 | 5.03 ± 0.03 | 9.76 ± 0.22 | 14.64 ± 0.50 | |
| | | | 6.38 ± 0.09 | | | |
| Day 10 2 | 2.89 ± 0.07 | 4.31 ± 0.12 | 7.08 ± 0.15 | | | |
| Day 10 3 | | 4.11 ± 0.04 | 5.96 ± 0.11 | 8.12 ± 0.11 | | |
| Day 10 4 | 3.19 ± 0.11 | 4.81 ± 0.05 | | | 8.96 ± 0.18 | |
| IL-10 Healthy 2 | | | | 11.19 ± 0.11 | | |
| | | | | 14.20 ± 0.17 | | |
| IL-10 Healthy 3 | | | 8.17 ± 0.10 | 10.13 ± 0.12 | 29.66 ± 0.51 | |
| | | | | 13.62 ± 0.24 | | |
| IL-10 Healthy 4 | | | 6.52 ± 0.08 | 9.77 ± 0.06 | | |
| IL-10 III 1 | | 3.20 ± 0.04 | | | | |
| | | 4.23 ± 0.05 | | | | |
| IL-10 III 2 | 1.78 ± 0.02 | 3.07 ± 0.04 | | | | |
| | 2.23 ± 0.02 | | | | | |
| IL-10 III 3 | 2.33 ± 0.03 | 2.91 ± 0.03 | | | | |

**Table S3.**
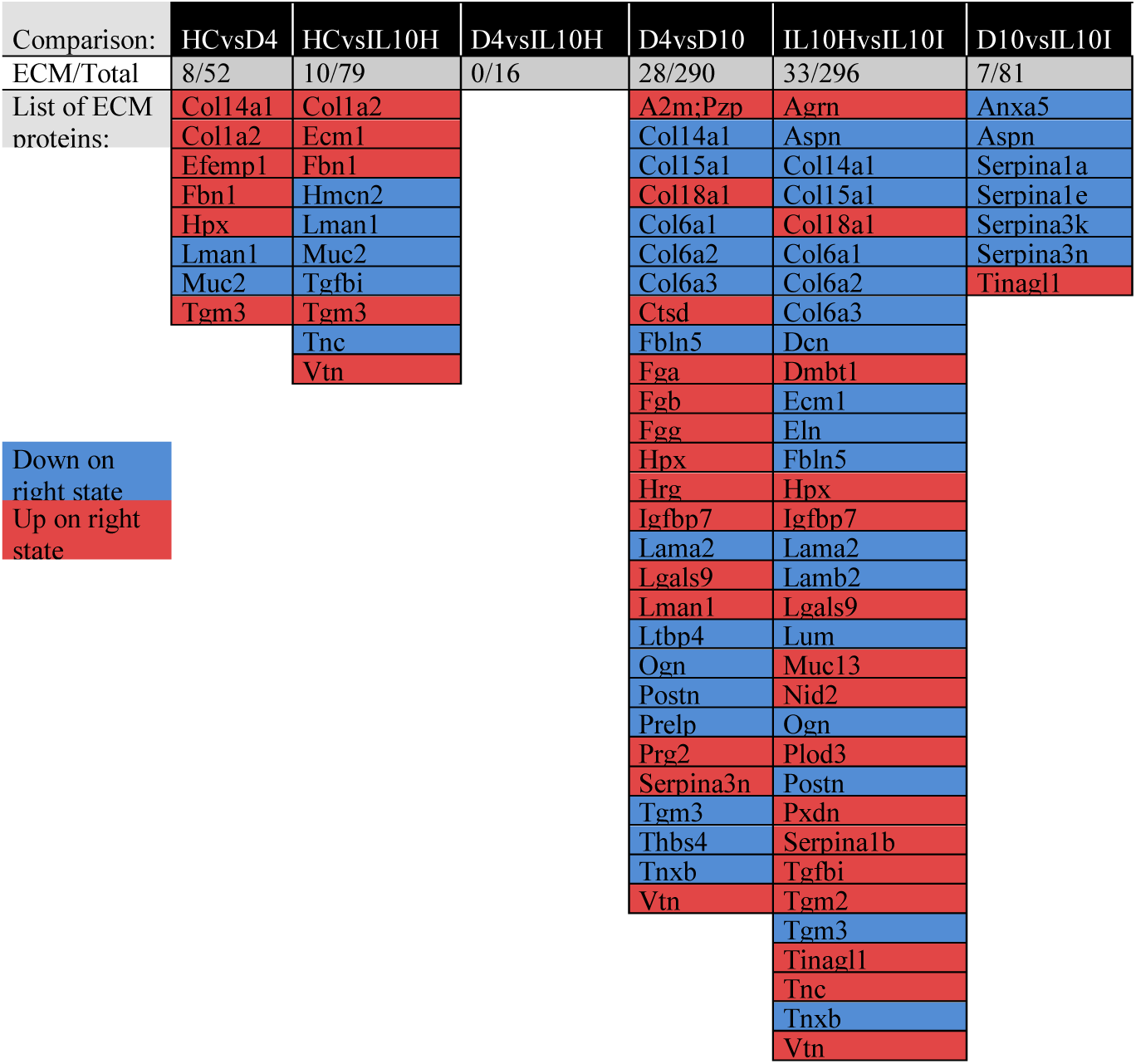
List of differentially abundant ECM proteins arising from MS results in comparisons of different state pairs. All differentially abundant proteins were identified by applying a t-test with a p-value cut off of 0.05 and permutation-based False Discovery Rate (FDR) correction. The second row indicates the proportion of ECM proteins out of the total number of differentially abundant proteins. Red indicates proteins that are more abundant on the state written on the right, and blue indicates proteins that are less abundant on the state written on the right. Abbreviations: HC=Healthy WT; D4=Day 4 of DSS model; D10=Day 4 of DSS model; IL10H=Healthy IL-10 mice; IL10I=Chronically ill IL-10 mice.

**Table S4.**
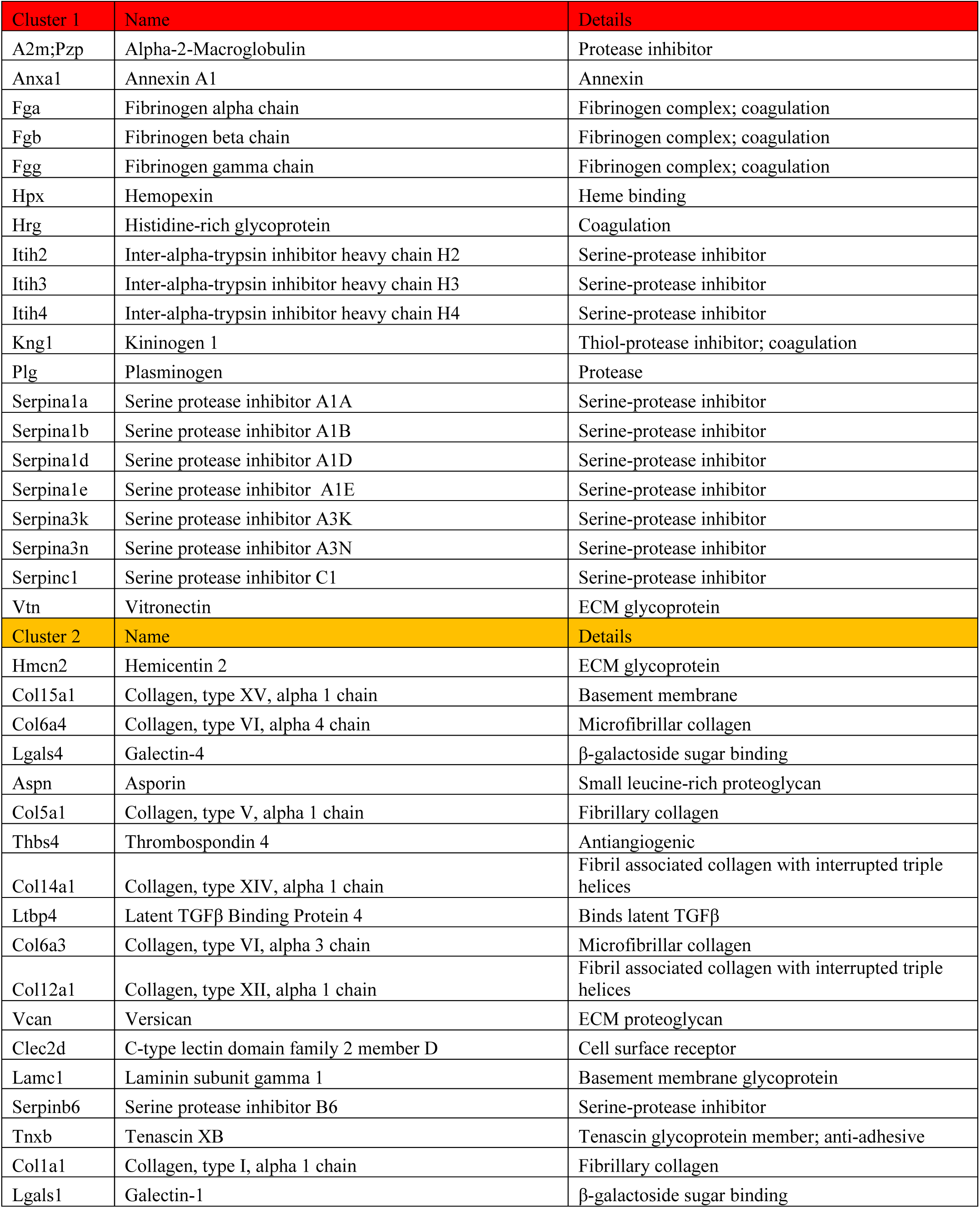
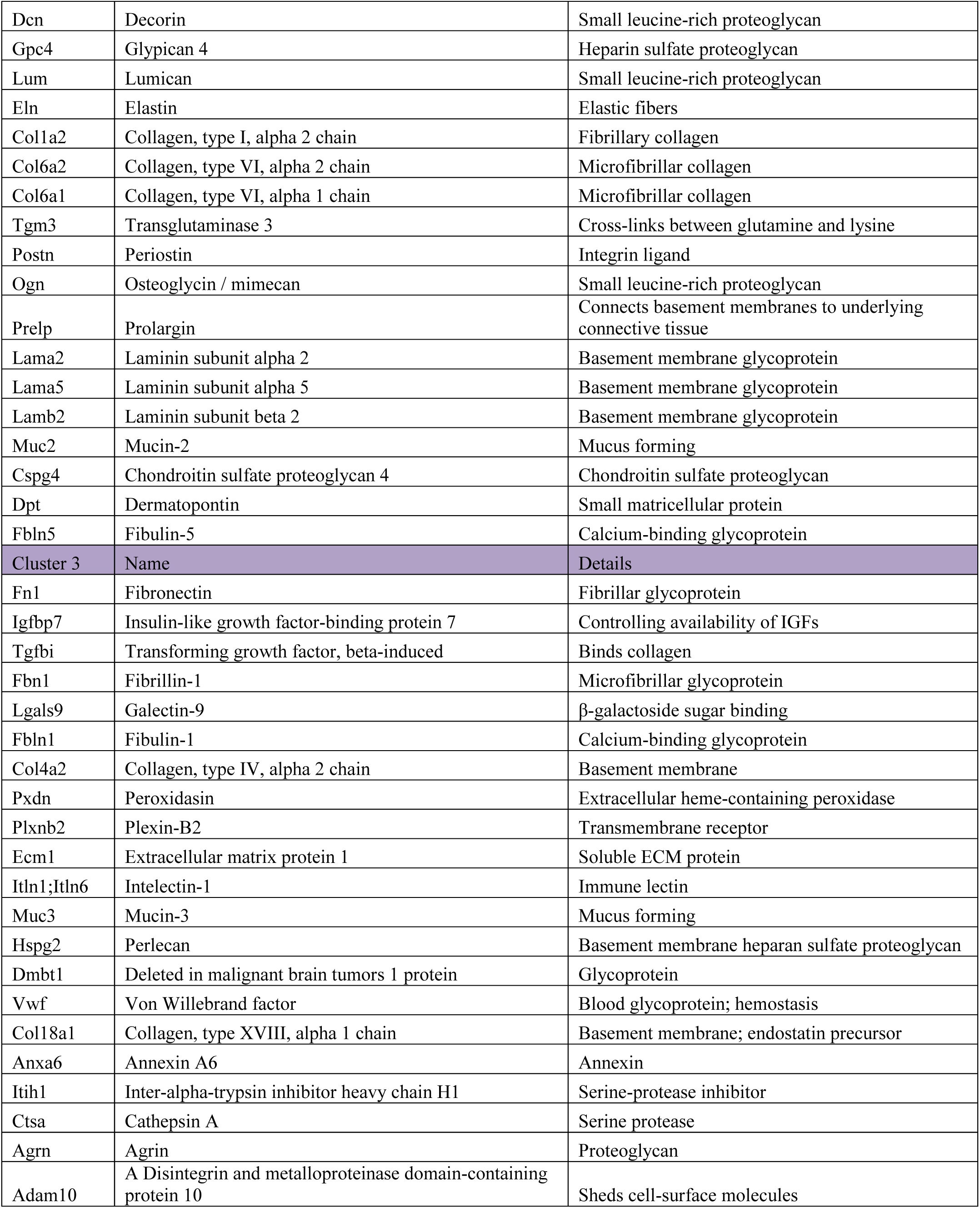
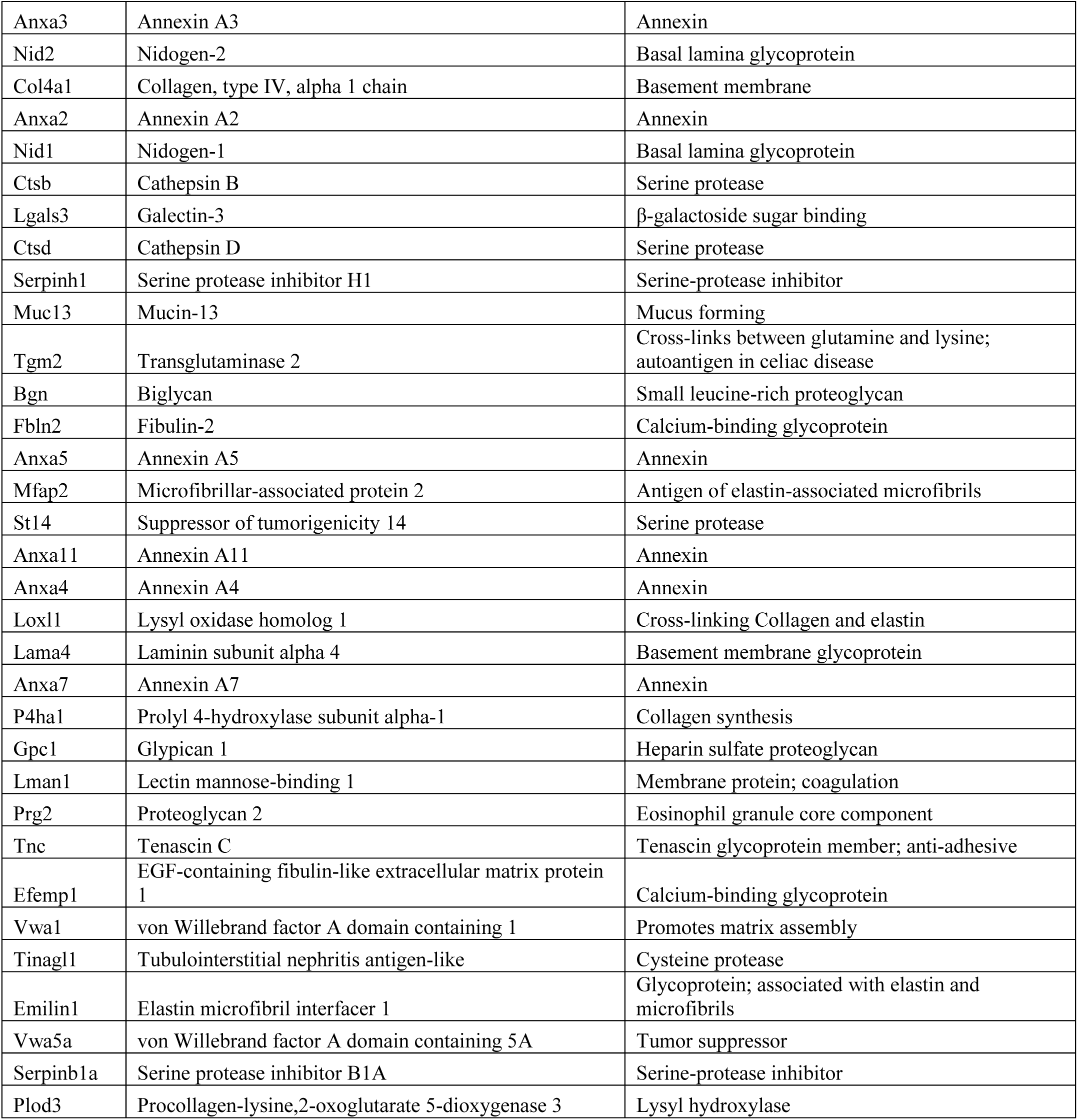
List of proteins in the three different clusters from **Fig. 3D** according to their abundance pattern across samples.

**Table S5.**
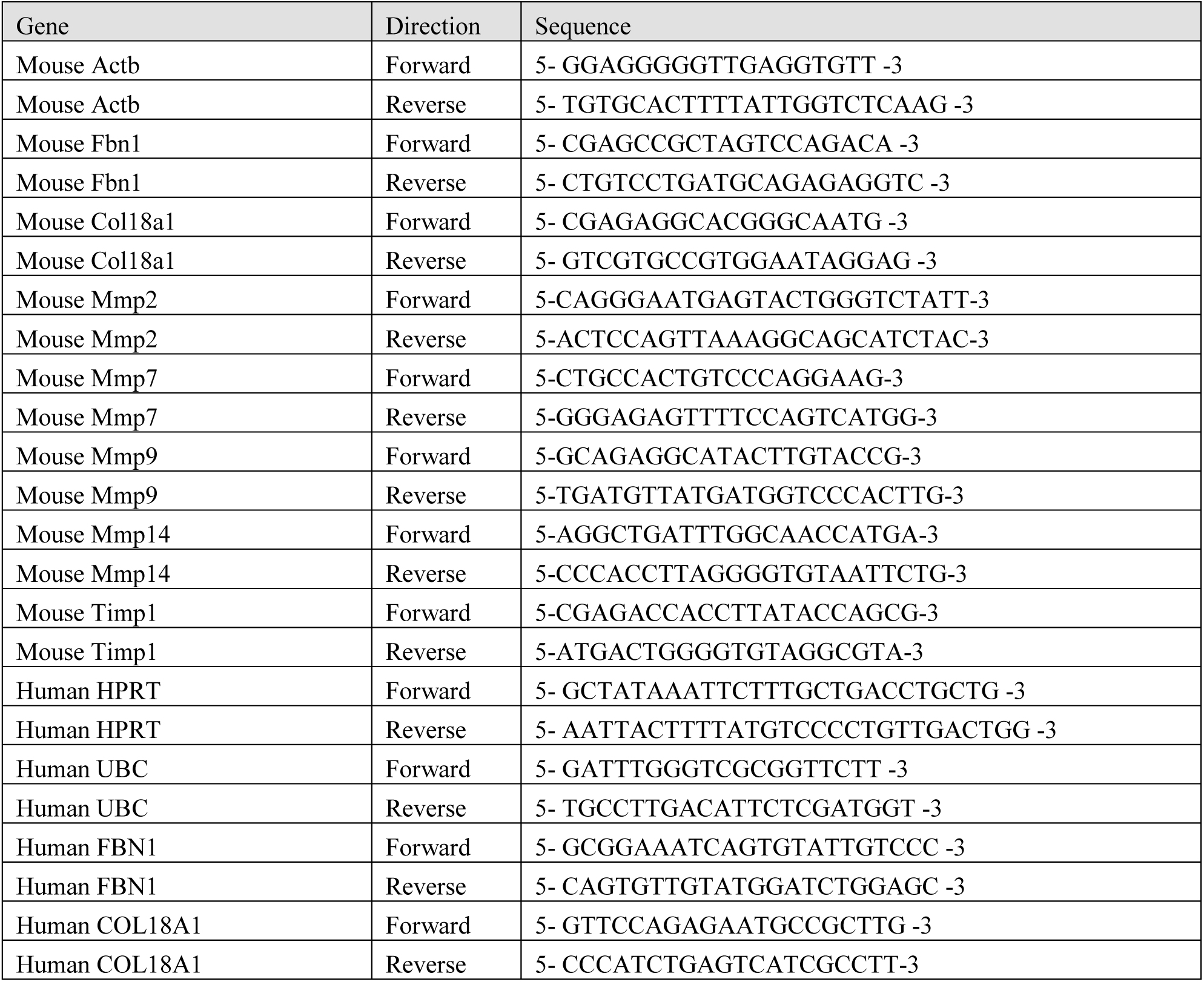
Primer sequences used for quantitative real-time PCR analysis.

**Table S6.** List of human samples in each group of mRNA analysis displayed in **Fig. 4F**.

| Sample type | Group | Type of IBD | Sex | Age |
| --- | --- | --- | --- | --- |
| Biopsy | Healthy | --- | M | 29 |
| Biopsy | Healthy | --- | M | 74 |
| Biopsy | Healthy | --- | F | 82 |
| Biopsy | Healthy | --- | F | 55 |
| Biopsy | Healthy | --- | F | 60 |
| Biopsy | Healthy | --- | M | 45 |
| Biopsy | Healthy | --- | F | 54 |
| Biopsy | Healthy | --- | F | 67 |
| Biopsy | Healthy | --- | M | 45 |
| Biopsy | Healthy | --- | M | 50 |
| Biopsy | Healthy | --- | M | 69 |
| Surgical resection | IBD | UC | F | 16 |
| Surgical resection | IBD | Undetermined | M | 43 |
| Surgical resection | IBD | UC | F | 36 |
| Surgical resection | IBD | Crohn | M | 48 |
| Surgical resection | IBD | Crohn | M | 38 |
| Surgical resection | IBD | Undetermined | F | 39 |
| Biopsy | IBD | UC | F | 26 |
| Biopsy | IBD | UC | F | 26 |
| Biopsy | IBD | UC | M | 53 |
| Biopsy | IBD | UC | M | 25 |

**Table S7.** List of antibodies used in the mass-cytometry analysis.

| Name | Product | Provider | Isotope conjugation |
| --- | --- | --- | --- |
| Anti-Human/ Mouse/ Rat Ki-67 | 3168007 | FLUIDIGM | 168Er - FLUIDIGM |
| Anti-Human/Mouse CD45R/B220 | 3160012 | FLUIDIGM | 160Gd – FLUIDIGM |
| Anti-Mouse CD11c | 3142003 | FLUIDIGM | 142Nd – FLUIDIGM |
| Anti-Mouse NK1.1 | 3170002 | FLUIDIGM | 170Er – FLUIDIGM |
| Anti-Mouse CD4 | 3145002 | FLUIDIGM | 145Nd – FLUIDIGM |
| Anti-Mouse CD8a | 3146003 | FLUIDIGM | 146Nd – FLUIDIGM |
| Anti-Mouse TER-119 | 3154005 | FLUIDIGM | 154Sm – FLUIDIGM |
| Anti-Mouse Ly-6C | 3162014 | FLUIDIGM | 162Dy – FLUIDIGM |
| Anti-Mouse Ly-6G | 3141008B | FLUIDIGM | 141Pr – FLUIDIGM |
| Anti-Mouse CD64 | 3151012 | FLUIDIGM | 151Eu – FLUIDIGM |
| Anti- Mouse F4/80 (BM8) | 3159009 | FLUIDIGM | 159Tb – FLUIDIGM |
| Anti-Mouse CD31/PECAM-1 (390) | 3165013 | FLUIDIGM | 165Ho – FLUIDIGM |
| Anti- Mouse CD326 [EpCAM] (G8.8) | 3166014 | FLUIDIGM | 166Er – FLUIDIGM |
| Anti-Mouse CD45 | 3089005 | FLUIDIGM | 89Y – FLUIDIGM |
| Anti-mouse CD3e (maxpar ready) | BLG-100345 | Biolegend | 153Eu – FLUIDIGM |
| Anti-Mouse CX3CR1 (SA011F11) | 3164023 | FLUIDIGM | 164Dy – FLUIDIGM |
| Anti-Mouse CD206 | 3169021B | FLUIDIGM | 169Tm – FLUIDIGM |
| Anti-Mouse I-A/I-E (M5/114.15.2) | 3209006B | FLUIDIGM | 209Bi – FLUIDIGM |
| Anti-Mouse CD11b | 3143015 | FLUIDIGM | 143Nd – FLUIDIGM |
| Anti-MMP9 | ab38898 | Abcam | 149Sm – Home made |
| Anti-MMP14 | ab51074 | Abcam | 156Gd – Home made |
| Anti-TIMP2 | ab38973 | Abcam | 172Yb – Home made |
| Anti-MMP2 | ab92536 | Abcam | 176Yb – Home made |
| Anti-ADAM-17 | ab57484 | Abcam | 174Yb – Home made |
| Anti-MMP7 | ab5706 | Abcam | 148Nd – Home made |
| Anti-CD34 | 14-0341-82 | Invitrogen | 152Sm – Home made |
| Anti-LOX | ab31238 | Abcam | 175Lu – Home made |
| Anti-αSMA | ab32575 | Abcam | 173Yb – Home made |
| Anti-MMP-13 | ab39012 | Abcam | 150Nd – Home made |

**Table S8.** Gating and cell population definition in the mass cytometry analysis.

| ID | Markers |
| --- | --- |
| Epithelial cells | CD326+ ; CD45- |
| Early endothelial progenitors | CD326- ; CD45- ; CD34+ ; CD31- |
| Late endothelial progenitors | CD326- ; CD45- ; CD34+ ; CD31+ |
| Adult Endothelial cells | CD326- ; CD45- ; CD34- ; CD31+ |
| Cytotoxic T lymphocytes | CD326- ; CD45+ ; CD11b- ; CD3e+ ; CD8a+ ; CD4- |
| T helper lymphocytes | CD326- ; CD45+ ; CD11b- ; CD3e+ ; CD8a- ; CD4+ |
| B cells | CD326- ; CD45+ ; CD11b- ; CD3e- ; CD45R+ |
| Dendritic cells | CD326- ; CD45+ ; CD11c+ ; CD3e- ; CD45R- ; MHC class II + |
| Neutrophils | CD326- ; CD45+ ; CD11b+ ; CD3e- ; CD45R- ; Ly-6G+ |
| Macrophages | CD326- ; CD45+ ; CD11b+ ; CD3e- ; CD45R- ; Ly-6G- ; F4/80+ |
| Circulating (pro inflammatory) monocytes | CD326- ; CD45+ ; CD11b- ; CD3e- ; CD45R- ; Ly-6G- ; F4/80- ; Ly-6Chi |

## References

[1] M. Grossman, N. Ben-Chetrit, A. Zhuravlev, R. Afik, E. Bassat, I. Solomonov, Y. Yarden, I. Sagi, Tumor cell invasion can be blocked by modulators of collagen fibril alignment that control assembly of the extracellular matrix, Cancer Res. 76 (2016) 4249–4258. https://doi.org/10.1158/0008-5472.CAN-15-2813.

[2] J. Herrera, C.A. Henke, P.B. Bitterman, Extracellular matrix as a driver of progressive fibrosis, J. Clin. Invest. 128 (2018) 45–53. https://doi.org/10.1172/JCI93557.

[3] P. Lu, V.M. Weaver, Z. Werb, The extracellular matrix: A dynamic niche in cancer progression, J. Cell Biol. 196 (2012) 395–406. https://doi.org/10.1083/jcb.201102147.

[4] R. Khokha, A. Murthy, A. Weiss, Metalloproteinases and their natural inhibitors in inflammation and immunity, Nat. Rev. Immunol. 13 (2013) 649–665. https://doi.org/10.1038/nri3499.

[5] E. Shimshoni, D. Yablecovitch, L. Baram, I. Dotan, I. Sagi, ECM remodelling in IBD: innocent bystander or partner in crime? The emerging role of extracellular molecular events in sustaining intestinal inflammation, Gut. 64 (2015) 367–72. https://doi.org/10.1136/gutjnl-2014-308048.

[6] M.D. Baugh, M.J. Perry, A.P. Hollander, D.R. Davies, S.S. Cross, A.J. Lobo, C.J. Taylor, G.S. Evans, Matrix metalloproteinase levels are elevated in inflammatory bowel disease., Gastroenterology. 117 (1999) 814–822.

[7] M.J.W. Meijer, M.A.C. Mieremet-Ooms, A.M. van der Zon, W. van Duijn, R.A. van Hogezand, C.F.M. Sier, D.W. Hommes, C.B.H.W. Lamers, H.W. Verspaget, Increased mucosal matrix metalloproteinase-1, −2, −3 and −9 activity in patients with inflammatory bowel disease and the relation with Crohn’s disease phenotype., Dig. Liver Dis. 39 (2007) 733–9. https://doi.org/10.1016/j.dld.2007.05.010.

[8] G. Pedersen, T. Saermark, T. Kirkegaard, J. Brynskov, Spontaneous and cytokine induced expression and activity of matrix metalloproteinases in human colonic epithelium, Clin. Exp. Immunol. 155 (2009) 257–265. https://doi.org/10.1111/j.1365-2249.2008.03836.x.

[9] F.E. Castaneda, B. Walia, M. Vijay-Kumar, N.R. Patel, S. Roser, V.L. Kolachala, M. Rojas, L. Wang, G. Oprea, P. Garg, A.T. Gewirtz, J. Roman, D. Merlin, S. V Sitaraman, Targeted deletion of metalloproteinase 9 attenuates experimental colitis in mice: central role of epithelial-derived MMP., Gastroenterology. 129 (2005) 1991–2008. https://doi.org/10.1053/j.gastro.2005.09.017.

[10] P. Garg, M. Vijay-Kumar, L. Wang, A.T. Gewirtz, D. Merlin, S. V Sitaraman, Matrix metalloproteinase-9-mediated tissue injury overrides the protective effect of matrix metalloproteinase-2 during colitis., Am. J. Physiol. Gastrointest. Liver Physiol. 296 (2009) G175–G184. https://doi.org/10.1152/ajpgi.90454.2008.

[11] N. Sela-Passwell, R. Kikkeri, O. Dym, H. Rozenberg, R. Margalit, R. Arad-Yellin, M. Eisenstein, O. Brenner, T. Shoham, T. Danon, A. Shanzer, I. Sagi, Antibodies targeting the catalytic zinc complex of activated matrix metalloproteinases show therapeutic potential., Nat. Med. 18 (2012) 143–7. https://doi.org/10.1038/nm.2582.

[12] K. Lohr, H. Sardana, S. Lee, F. Wu, D.L. Huso, A.R. Hamad, S. Chakravarti, Extracellular matrix protein lumican regulates inflammation in a mouse model of colitis, Inflamm. Bowel Dis. 18 (2012) 143–151. https://doi.org/10.1002/ibd.21713.Extracellular.

[13] F.L. Koller, E.A. Dozier, K.T. Nam, M. Swee, T.P. Birkland, W.C. Parks, B. Fingleton, Lack of MMP10 exacerbates experimental colitis and promotes development of inflammation-associated colonic dysplasia., Lab. Invest. 92 (2012) 1749–59. https://doi.org/10.1038/labinvest.2012.141.

[14] M. Waterman, O. Ben-Izhak, R. Eliakim, G. Groisman, I. Vlodavsky, N. Ilan, Heparanase upregulation by colonic epithelium in inflammatory bowel disease., Mod. Pathol. 20 (2007) 8–14. https://doi.org/10.1038/modpathol.3800710.

[15] I. Okayasu, S. Hatakeyama, M. Yamada, T. Ohkusa, Y. Inagaki, R. Nakaya, A novel method in the induction of reliable experimental acute and chronic ulcerative colitis in mice., Gastroenterology. 98 (1990) 694–702.

[16] D. Berg, J. Zhang, J. Weinstock, H. Ismail, Rapid development of colitis in NSAID-treated IL-10–deficient mice, Gastroenterology. (2002) 1527–1542. https://doi.org/10.1053/gast.rostaglandins.

[17] E.F. Stange, S.P.L. Travis, S. Vermeire, C. Beglinger, L. Kupcinkas, K. Geboes, A. Barakauskiene, V. Villanacci, A. Von Herbay, B.F. Warren, C. Gasche, H. Tilg, S.W. Schreiber, J. Schölmerich, W. Reinisch, European Crohn’s and Colitis Organisation, European evidence based consensus on the diagnosis and management of Crohn’s disease: definitions and diagnosis., Gut. 55 Suppl 1 (2006) i1–15. https://doi.org/10.1136/gut.2005.081950a.

[18] R. Kühn, J. Löhler, D. Rennick, K. Rajewsky, W. Müller, Interleukin-10-deficient mice develop chronic enterocolitis., Cell. 75 (1993) 263–274. https://doi.org/10.1016/0092-8674(93)80068-P.

[19] E.-O. Glocker, D. Kotlarz, K. Boztug, E.M. Gertz, A.A. Schäffer, F. Noyan, M. Perro, J. Diestelhorst, A. Allroth, D. Murugan, N. Hätscher, D. Pfeifer, K.-W. Sykora, M. Sauer, H. Kreipe, M. Lacher, R. Nustede, C. Woellner, U. Baumann, U. Salzer, S. Koletzko, N. Shah, A.W. Segal, A. Sauerbrey, S. Buderus, S.B. Snapper, B. Grimbacher, C. Klein, Inflammatory bowel disease and mutations affecting the interleukin-10 receptor., N. Engl. J. Med. 361 (2009) 2033–45. https://doi.org/10.1056/NEJMoa0907206.

[20] M. van der Rest, R. Garrone, Collagen family of proteins., FASEB J. (1991).

[21] M. Indrieri, A. Podestà, G. Bongiorno, D. Marchesi, P. Milani, Adhesive-free colloidal probes for nanoscale force measurements: production and characterization., Rev. Sci. Instrum. 82 (2011) 023708. https://doi.org/10.1063/1.3553499.

[22] L. Puricelli, M. Galluzzi, C. Schulte, A. Podestà, P. Milani, Nanomechanical and topographical imaging of living cells by atomic force microscopy with colloidal probes., Rev. Sci. Instrum. 86 (2015) 033705. https://doi.org/10.1063/1.4915896.

[23] A. Naba, K.R. Clauser, S. Hoersch, H. Liu, S.A. Carr, R.O. Hynes, The Matrisome: In Silico Definition and In Vivo Characterization by Proteomics of Normal and Tumor Extracellular Matrices, Mol. Cell. Proteomics. 11 (2012) M111.014647-M111.014647. https://doi.org/10.1074/mcp.M111.014647.

[24] T.B. Bennike, T.G. Carlsen, T. Ellingsen, O.K. Bonderup, H. Glerup, M. Bøgsted, G. Christiansen, S. Birkelund, A. Stensballe, V. Andersen, Neutrophil Extracellular Traps in Ulcerative Colitis: A Proteome Analysis of Intestinal Biopsies., Inflamm. Bowel Dis. 21 (2015) 2052–67. https://doi.org/10.1097/MIB.0000000000000460.

[25] A.G. Marneros, B.R. Olsen, Physiological role of collagen XVIII and endostatin., FASEB J. 19 (2005) 716–28. https://doi.org/10.1096/fj.04-2134rev.

[26] A. Utriainen, R. Sormunen, M. Kettunen, L.S. Carvalhaes, E. Sajanti, L. Eklund, R. Kauppinen, G.T. Kitten, T. Pihlajaniemi, Structurally altered basement membranes and hydrocephalus in a type XVIII collagen deficient mouse line, Hum. Mol. Genet. 13 (2004) 2089–2099. https://doi.org/10.1093/hmg/ddh213.

[27] L.Y. Sakai, D.R. Keene, E. Engvall, Fibrillin, a new 350-kD glycoprotein, is a component of extracellular microfibrils., J. Cell Biol. 103 (1986) 2499–509. http://www.ncbi.nlm.nih.gov/pubmed/3536967 (accessed July 20, 2017).

[28] P.E. Van den Steen, B. Dubois, I. Nelissen, P.M. Rudd, R.A. Dwek, G. Opdenakker, Biochemistry and Molecular Biology of Gelatinase B or Matrix Metalloproteinase-9 (MMP-9), Crit. Rev. Biochem. Mol. Biol. 37 (2002) 375–536. https://doi.org/10.1080/10409230290771546.

[29] Z.-S. Zeng, A.M. Cohen, J.G. Guillem, Loss of basement membrane type IV collagen is associated with increased expression of metalloproteinases 2 and 9 (MMP-2 and MMP-9) during human colorectal tumorigenesis, Carcinogenesis. 20 (1999) 749–755. https://doi.org/10.1093/carcin/20.5.749.

[30] J. Gaffney, I. Solomonov, E. Zehorai, I. Sagi, Multilevel regulation of matrix metalloproteinases in tissue homeostasis indicates their molecular specificity in vivo, Matrix Biol. 44–46 (2015) 191–199. https://doi.org/10.1016/J.MATBIO.2015.01.012.

[31] S.J. George, J.L. Johnson, In Situ Zymography, in: Humana Press, Totowa, NJ, 2010: pp. 271–277. https://doi.org/10.1007/978-1-60327-299-5_17.

[32] A. Kofla-Dlubacz, M. Matusiewicz, M. Krzystek-Korpacka, B. Iwanczak, Correlation of MMP-3 and MMP-9 with crohn’s disease activity in children, Dig. Dis. Sci. 57 (2012) 706–712. https://doi.org/10.1007/s10620-011-1936-z.

[33] A.M. Howard, K.S. Lafever, A.M. Fenix, C.R. Scurrah, K.S. Lau, D.T. Burnette, G. Bhave, N. Ferrell, A. Page-Mccaw, DSS-induced damage to basement membranes is repaired by matrix replacement and crosslinking, J. Cell Sci. 132 (2019). https://doi.org/10.1242/jcs.226860.

[34] S. Kugathasan, L.A. Denson, T.D. Walters, M.-O. Kim, U.M. Marigorta, M. Schirmer, K. Mondal, C. Liu, A. Griffiths, J.D. Noe, W. V Crandall, S. Snapper, S. Rabizadeh, J.R. Rosh, J.M. Shapiro, S. Guthery, D.R. Mack, R. Kellermayer, M.D. Kappelman, S. Steiner, D.E. Moulton, D. Keljo, S. Cohen, M. Oliva-Hemker, M.B. Heyman, A.R. Otley, S.S. Baker, J.S. Evans, B.S. Kirschner, A.S. Patel, D. Ziring, B.C. Trapnell, F.A. Sylvester, M.C. Stephens, R.N. Baldassano, J.F. Markowitz, J. Cho, R.J. Xavier, C. Huttenhower, B.J. Aronow, G. Gibson, J.S. Hyams, M.C. Dubinsky, Prediction of complicated disease course for children newly diagnosed with Crohn’s disease: a multicentre inception cohort study, Lancet. 389 (2017) 1710–1718. https://doi.org/10.1016/S0140-6736(17)30317-3.

[35] F. Rieder, J.R. de Bruyn, B.T. Pham, K. Katsanos, V. Annese, P.D.R. Higgins, F. Magro, I. Dotan, Results of the 4th Scientific Workshop of the ECCO (Group II): Markers of intestinal fibrosis in inflammatory bowel disease., J. Crohns. Colitis. (2014). https://doi.org/10.1016/j.crohns.2014.03.009.

[36] O.H. Nielsen, G. Rogler, D. Hahnloser, O.Ø. Thomsen, Diagnosis and management of fistulizing Crohn’s disease., Nat. Clin. Pract. Gastroenterol. Hepatol. 6 (2009) 92–106. https://doi.org/10.1038/ncpgasthep1340.

[37] G. Latella, G. Rogler, G. Bamias, C. Breynaert, J. Florholmen, G. Pellino, S. Reif, S. Speca, I.C. Lawrance, Results of the 4th scientific workshop of the ECCO (I): Pathophysiology of intestinal fibrosis in IBD., J. Crohns. Colitis. (2014). https://doi.org/10.1016/j.crohns.2014.03.008.

[38] F. Rieder, C. Fiocchi, Intestinal fibrosis in IBD — a dynamic, multifactorial process, 6 (2009) 228–235. https://doi.org/10.1038/nrgastro.2009.31.

[39] B.W. Bardoel, E.F. Kenny, G. Sollberger, A. Zychlinsky, The Balancing Act of Neutrophils, Cell Host Microbe. 15 (2014) 526–536. https://doi.org/10.1016/J.CHOM.2014.04.011.

[40] D. Talmi-Frank, Z. Altboum, I. Solomonov, Y. Udi, D.A. Jaitin, M. Klepfish, E. David, A. Zhuravlev, H. Keren-Shaul, D.R. Winter, I. Gat-Viks, M. Mandelboim, T. Ziv, I. Amit, I. Sagi, Extracellular Matrix Proteolysis by MT1-MMP Contributes to Influenza-Related Tissue Damage and Mortality., Cell Host Microbe. 20 (2016) 458–470. https://doi.org/10.1016/j.chom.2016.09.005.

[41] C. Becker, M.C. Fantini, M.F. Neurath, High resolution colonoscopy in live mice., Nat. Protoc. 1 (2006) 2900–2904. https://doi.org/10.1038/nprot.2006.446.

[42] E. Zigmond, C. Varol, J. Farache, E. Elmaliah, A.T. Satpathy, G. Friedlander, M. Mack, N. Shpigel, I.G. Boneca, K.M. Murphy, G. Shakhar, Z. Halpern, S. Jung, Ly6C hi Monocytes in the Inflamed Colon Give Rise to Proinflammatory Effector Cells and Migratory Antigen-Presenting Cells, Immunity. (2012). https://doi.org/10.1016/j.immuni.2012.08.026.

[43] G.K. Behbehani, C. Thom, E.R. Zunder, R. Finck, B. Gaudilliere, G.K. Fragiadakis, W.J. Fantl, G.P. Nolan, Transient partial permeabilization with saponin enables cellular barcoding prior to surface marker staining, Cytometry. A. 85 (2014) 1011. https://doi.org/10.1002/CYTO.A.22573.

[44] E.R. Zunder, R. Finck, G.K. Behbehani, E.D. Amir, S. Krishnaswamy, V.D. Gonzalez, C.G. Lorang, Z. Bjornson, M.H. Spitzer, B. Bodenmiller, W.J. Fantl, D. Pe’er, G.P. Nolan, Palladium-based mass tag cell barcoding with a doublet-filtering scheme and single-cell deconvolution algorithm, Nat. Protoc. 10 (2015) 316–333. https://doi.org/10.1038/nprot.2015.020.

[45] H. Lu, T. Hoshiba, N. Kawazoe, G. Chen, Comparison of decellularization techniques for preparation of extracellular matrix scaffolds derived from three-dimensional cell culture., J. Biomed. Mater. Res. A. 100 (2012) 2507–16. https://doi.org/10.1002/jbm.a.34150.

[46] E. Shimshoni, I. Sagi, Sample Preparation of Extracellular Matrix of Murine Colons for Scanning Electron Microscopy, in: I. Sagi, N. Afratis (Eds.), Collagen. Methods Mol. Biol., Humana Press, New York, NY, 2019: pp. 129–133. https://doi.org/10.1007/978-1-4939-9095-5_9.

[47] H.-J. Butt, B. Cappella, M. Kappl, Force measurements with the atomic force microscope: Technique, interpretation and applications, Surf. Sci. Rep. 59 (2005) 1–152. https://doi.org/10.1016/j.surfrep.2005.08.003.

[48] H. Schillers, C. Rianna, J. Schäpe, T. Luque, H. Doschke, M. Wälte, J.J. Uriarte, N. Campillo, G.P.A. Michanetzis, J. Bobrowska, A. Dumitru, E.T. Herruzo, S. Bovio, P. Parot, M. Galluzzi, A. Podestà, L. Puricelli, S. Scheuring, Y. Missirlis, R. Garcia, M. Odorico, J.-M. Teulon, F. Lafont, M. Lekka, F. Rico, A. Rigato, J.-L. Pellequer, H. Oberleithner, D. Navajas, M. Radmacher, Standardized Nanomechanical Atomic Force Microscopy Procedure (SNAP) for Measuring Soft and Biological Samples, Sci. Rep. 7 (2017) 5117. https://doi.org/10.1038/s41598-017-05383-0.

[49] H. Cramér, Mathematical Methods of Statistics, Princet. Univ. Press. (1999).

[50] J. Cox, M. Mann, MaxQuant enables high peptide identification rates, individualized p.p.b.-range mass accuracies and proteome-wide protein quantification., Nat. Biotechnol. 26 (2008) 1367–1372. https://doi.org/10.1038/nbt.1511.

[51] S. Tyanova, T. Temu, J. Cox, The MaxQuant computational platform for mass spectrometry – based shotgun proteomics, Nat. Protoc. 11 (2016) 2301–2319. https://doi.org/10.1038/nprot.2016.136.

[52] J. Cox, N. Neuhauser, A. Michalski, R. a. Scheltema, J. V. Olsen, M. Mann, Andromeda: A peptide search engine integrated into the MaxQuant environment, J. Proteome Res. 10 (2011) 1794–1805. https://doi.org/10.1021/pr101065j.

[53] J. Cox, M.Y. Hein, C.A. Luber, I. Paron, N. Nagaraj, M. Mann, Accurate Proteome-wide Label-free Quantification by Delayed Normalization and Maximal Peptide Ratio Extraction, Termed MaxLFQ, Mol. Cell. Proteomics. 13 (2014) 2513–2526. https://doi.org/10.1074/mcp.M113.031591.

[54] S.L. Salzberg, C4.5: Programs for Machine Learning by J. Ross Quinlan. Morgan Kaufmann Publishers, Inc., 1993, Mach. Learn. 16 (1994) 235–240. https://doi.org/10.1007/BF00993309.

[55] K.J. Livak, T.D. Schmittgen, Analysis of relative gene expression data using real-time quantitative PCR and the 2(-Delta Delta Ct) Method, Methods. 25 (2001) 402–408. https://doi.org/10.1006/meth.2001.1262.

[56] C. Leys, C. Ley, O. Klein, P. Bernard, L. Licata, Detecting outliers : Do not use standard deviation around the mean, use absolute deviation around the median, J. Exp. Soc. Psychol. (2013) 4–6.

[57] S. Tyanova, T. Temu, P. Sinitcyn, A. Carlson, M.Y. Hein, T. Geiger, M. Mann, J. Cox, The Perseus computational platform for comprehensive analysis of (prote)omics data, Nat. Methods. 13 (2016) 731–740. https://doi.org/10.1038/nmeth.3901.

[58] V.G. Tusher, R. Tibshirani, G. Chu, Significance analysis of microarrays applied to the ionizing radiation response, Proc. Natl. Acad. Sci. 98 (2001) 5116–5121. https://doi.org/10.1073/pnas.091062498.

[59] I. Solomonov, E. Zehorai, D. Talmi-Frank, S.G. Wolf, A. Shainskaya, A. Zhuravlev, E. Kartvelishvily, R. Visse, Y. Levin, N. Kampf, D.A. Jaitin, E. David, I. Amit, H. Nagase, I. Sagi, Distinct biological events generated by ECM proteolysis by two homologous collagenases., Proc. Natl. Acad. Sci. U. S. A. 113 (2016) 10884–9. https://doi.org/10.1073/pnas.1519676113.

